# Longevity-promoting human gut *Bifidobacteria* strains require distinct cytoprotective pathways and share dependence on host lipid regulation

**DOI:** 10.1101/2025.10.17.682934

**Authors:** Yuxin Li, Gabriela Diaz-Tang, Shuo Han

## Abstract

Human gut *Bifidobacteria* are abundant during infancy and have been reported to be enriched in some exceptionally long-lived populations, yet whether their beneficial effects on lifespan and healthspan are broadly shared across the genus or restricted to specific strains remains unclear. Using an anaerobic bacteria–*Caenorhabditis elegans* platform with heat-killed diets, we systematically compared 11 human gut *Bifidobacteria* strains representing nine species. *B. infantis* ATCC 15697, *B. longum* NCC 2705, and *B. breve* DSMZ 20213 produced the largest lifespan extensions and improved multiple measures of physiological resilience, whereas the other strains produced smaller, neutral, or detrimental effects. Genetic analyses showed that these three strains required different combinations of conserved cytoprotective regulators, yet all depended on NHR-49, a lipid-regulating nuclear receptor functionally related to mammalian PPARα, for full lifespan extension. Consistent with this shared requirement, further analyses showed that *B. infantis* and *B. longum* also required FAT-7, an NHR-49-regulated delta-9 fatty acid desaturase, for full lifespan extension and oxidative stress protection. Untargeted lipidomics identified distinct but partially overlapping phosphatidylethanolamine, diacylglycerol, and triacylglycerol species that were enriched by these diets and reduced by *fat-7* RNAi. Bulk lipid extracts from either strain enhanced oxidative stress resistance when added to a standard *E. coli* diet, providing functional evidence that bacterial lipids contribute to protection. Together, these findings support a strain-selective model. Related commensal strains differ in their physiological effects and cytoprotective pathway requirements but share dependence on host lipid regulation. The findings also identify gut bacteria–host lipid interactions as a mechanistic axis linking microbial products to stress resilience and longevity.

**Author summary:** Gut bacteria are increasingly linked to healthy aging, but which strains are beneficial and how they act on the host remain poorly understood. Human gut *Bifidobacteria* are common members of the infant gut microbiome and have been reported to be enriched in some exceptionally long-lived populations. Using *Caenorhabditis elegans* and controlled, heat-killed bacterial diets, we compared 11 human gut *Bifidobacteria* strains representing nine species. *B. infantis*, *B. longum*, and *B. breve* produced the largest lifespan benefits and improved physiological resilience, whereas the remaining strains produced smaller, neutral, or detrimental effects, showing that these benefits are highly strain-selective. Genetic experiments revealed that the three selected strains required different combinations of conserved stress response regulators. Despite these differences, all depended on NHR-49, a regulator of host lipid metabolism functionally related to mammalian PPARα, for full lifespan extension. Consistent with this shared requirement, *B. infantis* and *B. longum* also required FAT-7, an NHR-49-regulated fatty acid desaturase, for lifespan extension and protection from oxidative stress. Lipidomic profiling showed that the two strains reshaped host complex lipid composition in partially overlapping ways, with many changes reduced when *fat-7* expression was lowered. Lipid extracts from either strain were also sufficient to improve oxidative stress resistance when added to a standard diet, providing functional evidence that bacterial lipids contribute to protection. Together, these findings identify gut bacteria–host lipid interactions as a mechanistic axis linking strain-specific bacterial effects to stress resilience and longevity.

## Introduction

The gut microbiome has emerged as an important modulator of host aging and longevity^1–4^, yet the mechanisms underlying these effects remain poorly resolved at the strain level. Among human gut bacteria, *Bifidobacteria* are abundant during infancy^5–7^ and have been reported to be enriched in some populations of exceptionally long-lived people^8,9^. Selected *Bifidobacteria* strains can extend lifespan or improve age-associated physiology in *Caenorhabditis elegans*^10–12^, *Drosophila*^12^, and mice^12,13^. These observations raise two unresolved questions. First, how extensively do lifespan and physiological benefits vary across *Bifidobacteria* strains? Second, do beneficial strains act through common host mechanisms, or do they engage distinct host pathways that converge on shared physiological outcomes?

To address these questions, we used *C. elegans* as a tractable system for controlled dietary comparisons and functional genetic analysis. Although nematodes and mammals differ greatly in biological complexity, *C. elegans* retains conserved stress response, metabolic, and longevity pathways^14,15^. The model enables systematic comparisons of bacterial diets^16,17^ and rapid genetic testing of host pathway requirements^18,19^ at a scale difficult to achieve in mammalian systems. Previous studies have identified lifespan effects of several bacterial taxa in *C. elegans*^10–12,20,21^ and linked individual *Bifidobacteria* strains to conserved regulators such as DAF-16/FOXO and SKN-1/Nrf2^10,11^. This combination of controlled dietary exposure and genetic tractability provides an experimental platform for comparing strain-level physiological effects and defining their host pathway requirements.

Here, we compared 11 human gut *Bifidobacteria* strains representing nine prevalent species^22,23^ and identified *B. longum* NCC 2705, *B. infantis* ATCC 15697, and *B. breve* DSMZ 20213 as the strains that produced the largest lifespan benefits and improved multiple measures of physiological resilience. These strains required different combinations of conserved cytoprotective regulators, yet all depended on NHR-49, a lipid-regulating nuclear receptor functionally related to mammalian PPARα. Further analyses of *B. infantis* and *B. longum* identified a shared dependence on FAT-7, revealed distinct but overlapping changes in host lipid composition, and implicated bacterial lipids as functional contributors to protection. Collectively, these findings show that strain-specific and shared mechanisms can coexist: related commensal strains can differ substantially in their effects and cytoprotective pathway requirements while sharing a common dependence on host lipid regulation.

### An anaerobic bacteria–*C. elegans* platform identifies three human gut *Bifidobacteria* strains with the largest lifespan benefits

To compare prevalent human gut *Bifidobacteria* strains under controlled conditions, we established an anaerobic bacteria-*C. elegans* platform (Fig 1A). Each strain was cultured anaerobically and heat-killed. We then normalized the bacterial biomass and provided each strain as a diet beginning on the first day of reproductive adulthood. Animals were maintained on live *E. coli* during development and then transferred to individual *Bifidobacteria* strains or a heat-killed *E. coli* control. This design reduced potential confounding from live bacterial growth, developmental exposure, and differences in bacterial dose.

**Fig 1.**
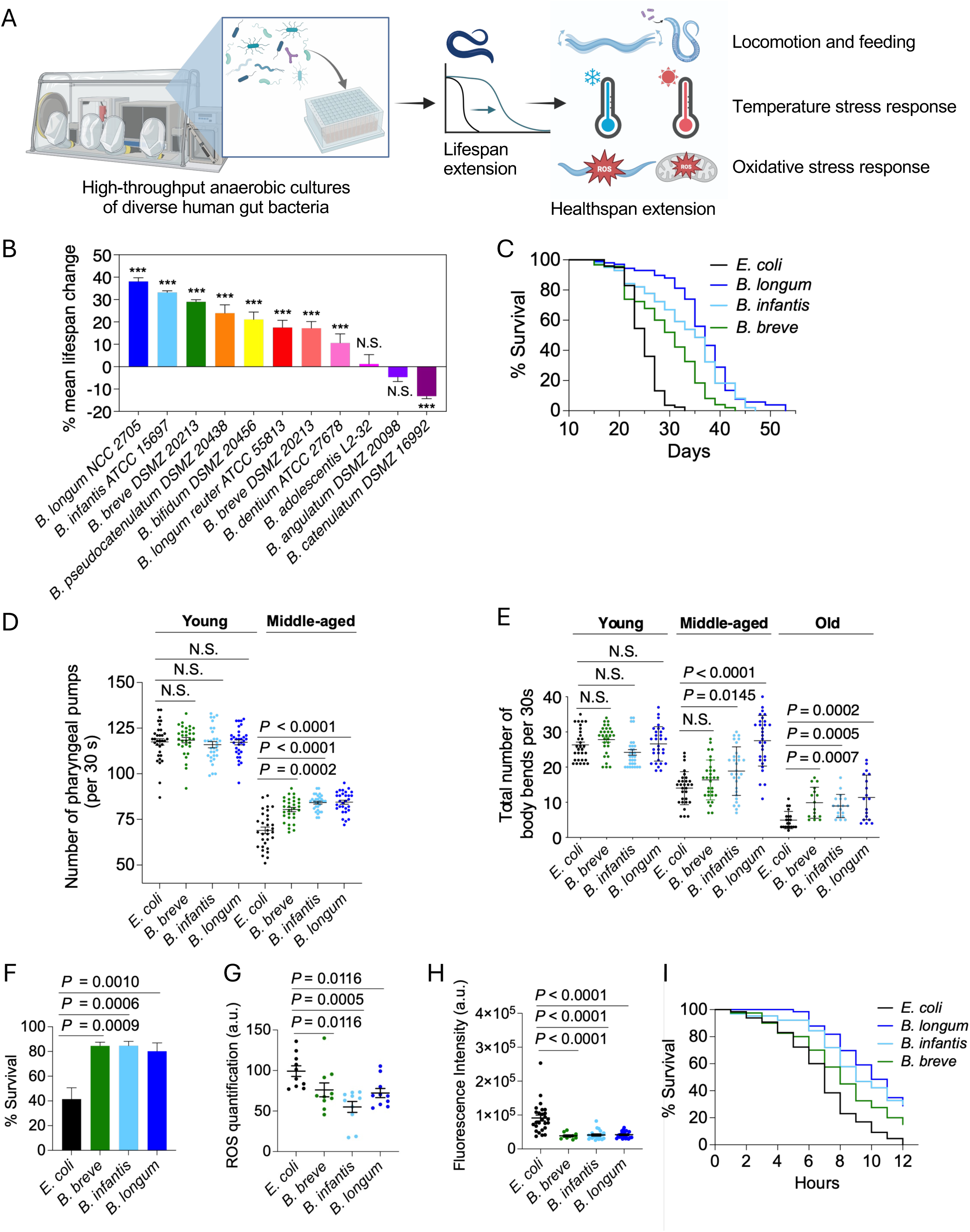
Distinct *Bifidobacteria* strains extend lifespan and aspects of healthspan. **A**. Schematic of the human anaerobic gut bacteria-*C. elegans* aging platform. Lifespan serves as the primary screen to identify top lifespan-extending strains, which are subsequently linked to distinct host healthspan metrics for downstream mechanistic studies. **B**. Lifespan screen across 11 *Bifidobacteria* strains reveals divergent lifespan responses. Bar graph shows mean lifespan change (%) of worms fed individual heat-killed *Bifidobacteria* strains comparing to those fed *E. coli* OP50 control. Distinct bacterial strains are introduced at day one of reproductive adulthood. Mean ± s.e.m. of six replicate plates per condition from two independent experiments, with three replicates per experiment; *n* > 30 per replicate. \*\*\**P* < 0.0001 by log-rank test; N.S., not significant. **C**. Example lifespan curves from the screen in **B** for worms fed each of the top three lifespan-extending *Bifidobacteria* strains (*B. longum* NCC 2705, *B. infantis* ATCC 15697, and *B. breve* DSMZ 20213, abbreviated as *B. longum*, *B. infantis*, and *B. breve* hereafter). **D**. Pharyngeal pumping rates of young (adult day 3) and middle-aged (adult day 8) worms. Pumping activity was quantified as the total number of pumps completed in 30 seconds per worm. Each dot represents one worm; *n* > 30 per condition. **E**. Locomotion of young (adult day 2), middle-aged (adult day 10), and old worms (adult day 20). Locomotion was quantified as the total number of body bends completed in 30 seconds per worm. Each dot represents one worm; *n* = 30 per condition for young and middle-aged worms, n ≥16 for old worms. **F**. Cold stress resistance of middle-aged worms. Survival (%) following exposure to 4°C for 60 hours; *n* > 30 per replicate plate. Mean ± s.e.m. of eleven replicate plates across two independent experiments. **G**. Reactive Oxygen Species (ROS) levels in middle-aged (adult day 8) worm lysates. Each dot represents an individual replicate plate with > 300 worms. Mean ± s.e.m. of nine replicate plates across three independent experiments, each with three biological replicates. **H**. MitoSOX fluorescence intensity in pharyngeal bulb muscle of live worms. Each dot represents one worm, n ≥ 12 per condition. Mean ± s.e.m.; representative of two independent experiments. **I**. Oxidative stress resistance of middle-aged (adult day 8) worms. Survival (%) following acute paraquat exposure (200 mM). Paraquat treatment was terminated within 12 hours. Survival statistics for all conditions were calculated at this endpoint. *n* ≥ 37 per condition. *P* ≤ 0.0031 by log-rank between individual *Bifidobacteria* strains and the *E. coli* control; representative of two independent experiments. **D**-**H***. P* values: Kruskal-Wallis with Benjamini Hochberg correction for multiple comparisons.

We tested 11 strains representing nine prevalent human gut *Bifidobacteria* species^22,23^ (S1 Table). Their effects on lifespan varied substantially. *B. longum* NCC 2705, *B. infantis* ATCC 15697, and *B. breve* DSMZ 20213 produced the largest lifespan extensions relative to the *E. coli* control (Fig 1B and 1C). The remaining strains produced smaller, neutral, or detrimental effects. In particular, *B. adolescentis* L2-32 and *B. angulatum* DSMZ 20098 did not significantly alter lifespan, whereas *B. catenulatum* DSMZ 16992 shortened lifespan (Fig 1B). These findings show that closely related members of the same bacterial genus can differ substantially in the magnitude and direction of their effects on host lifespan. We therefore selected the three strongest lifespan-extending strains for subsequent functional and mechanistic analyses and refer to them hereafter as *B. longum*, *B. infantis*, and *B. breve*.

To determine whether their lifespan effects persisted in the presence of the standard *E. coli* diet, we tested mixtures containing *E. coli* and each selected *Bifidobacteria* strain at 1:1 or 1:4 ratios while maintaining the same total bacterial biomass (S1A Fig). The 1:1 mixtures retained significant but attenuated lifespan benefits, whereas the 1:4 mixtures largely recapitulated the effects of the corresponding pure *Bifidobacteria* diets. This dose-dependent relationship between the proportion of *Bifidobacteria* and lifespan extension supports an active contribution of *Bifidobacteria* biomass or products to host longevity. The divergent lifespan responses across strains further indicate that this contribution is strain-specific rather than a general difference between *Bifidobacteria* and *E. coli* as dietary sources. Because mixed diets complicate attribution of bacterial products and host responses to individual strains, we used pure *Bifidobacteria* diets for subsequent mechanistic analyses.

### The three selected *Bifidobacteria* strains improve overlapping measures of physiological resilience

We next asked whether the three selected strains also influenced functional decline and stress resilience. We assessed pharyngeal pumping as a measure of feeding function, locomotion as a measure of age-associated mobility decline, and survival under cold, heat, and oxidative stress. Throughout this study, age groups were defined relative to animals fed the *E. coli* control diet. Young adults were sampled on days 2–3 of adulthood, middle-aged animals on days 8–10 as the control population approached midlife, and old animals on day 20 near the median lifespan of the control population.

Pharyngeal pumping rates did not differ among diets in young adults (Fig 1D). By middle age, however, all three *Bifidobacteria* strains increased pumping rates relative to the *E. coli* control, indicating that the diets did not impair food intake and were associated with preserved pharyngeal function (Fig 1D). Locomotor activity likewise did not differ in young adults. In middle-aged and old animals, *B. longum* produced the strongest improvement in locomotion, followed by *B. infantis* and *B. breve* (Fig 1E). Because these functional differences were most evident in middle-aged animals, we used this stage to examine stress resilience.

All three strains increased survival following cold exposure at 4°C (Fig 1F), whereas none improved survival under heat stress at 34°C (S1B Fig), showing that their protective effects were stressor-dependent. We next examined oxidative stress. All three strains reduced total reactive oxygen species in whole worm lysates and lowered mitochondrial superoxide in pharyngeal muscle (Fig 1G and 1H). We then asked whether these reductions were accompanied by improved survival under acute oxidative stress. Because paraquat is commonly used to induce ROS accumulation and oxidative stress responses in vivo^24^, we measured survival following paraquat exposure. *B. longum* and *B. infantis* significantly increased survival, whereas *B. breve* conferred a more modest benefit (Fig 1I). By contrast, two strains that did not extend lifespan, *B. angulatum* and *B. adolescentis*, did not protect against paraquat-induced oxidative stress (S1C Fig).

Together, these findings show that the three strains with the largest lifespan benefits also improved multiple measures of physiological resilience, although their effects on individual healthspan metrics differed. Across the three strains, those with larger lifespan benefits also tended to confer stronger oxidative stress protection. This correspondence suggests that overlapping mechanisms within each strain may contribute to both phenotypes, while shared components of these mechanisms may also operate across strains. We therefore asked whether these observations reflect shared and strain-specific requirements for conserved host pathways.

### The three selected *Bifidobacteria* strains share dependence on NHR-49 but differ in cytoprotective pathway requirements

Longevity in *C. elegans* is regulated by conserved stress response, metabolic, and lysosomal pathways^14,15^. We therefore asked whether the three strongest lifespan-extending *Bifidobacteria* strains required shared host regulators or distinct combinations of pathways. To test this, we performed a genetic interaction screen using a representative panel of established longevity regulators with functionally related mammalian counterparts: DAF-16/FOXO^25^, SKN-1/Nrf2^26^, HSF-1/HSF1^27,28^, ATFS-1/ATF5^29^, NHR-49/PPARα^30,31^, XBP-1/XBP1^32,33^, NHR-80/HNF4α^34^, SBP-1/SREBP1^35^, and HLH-30/TFEB^36^ (S2 Table). These factors were selected because they regulate worm longevity and are evolutionarily conserved, increasing the potential relevance of mechanisms identified here to mammalian hosts. We measured the lifespan benefit conferred by *B. infantis*, *B. longum*, or *B. breve* in wild-type animals and in animals deficient for each regulator by mutation or RNAi knockdown. We summarized the results in a heatmap in which rows show the average percentage of lifespan extension conferred by each strain relative to matched *E. coli*-fed controls across independent experiments, and columns represent wild-type or animals deficient in individual transcription factors through mutation or RNAi knockdown (Fig 2A and S3 Table).

**Fig 2.**
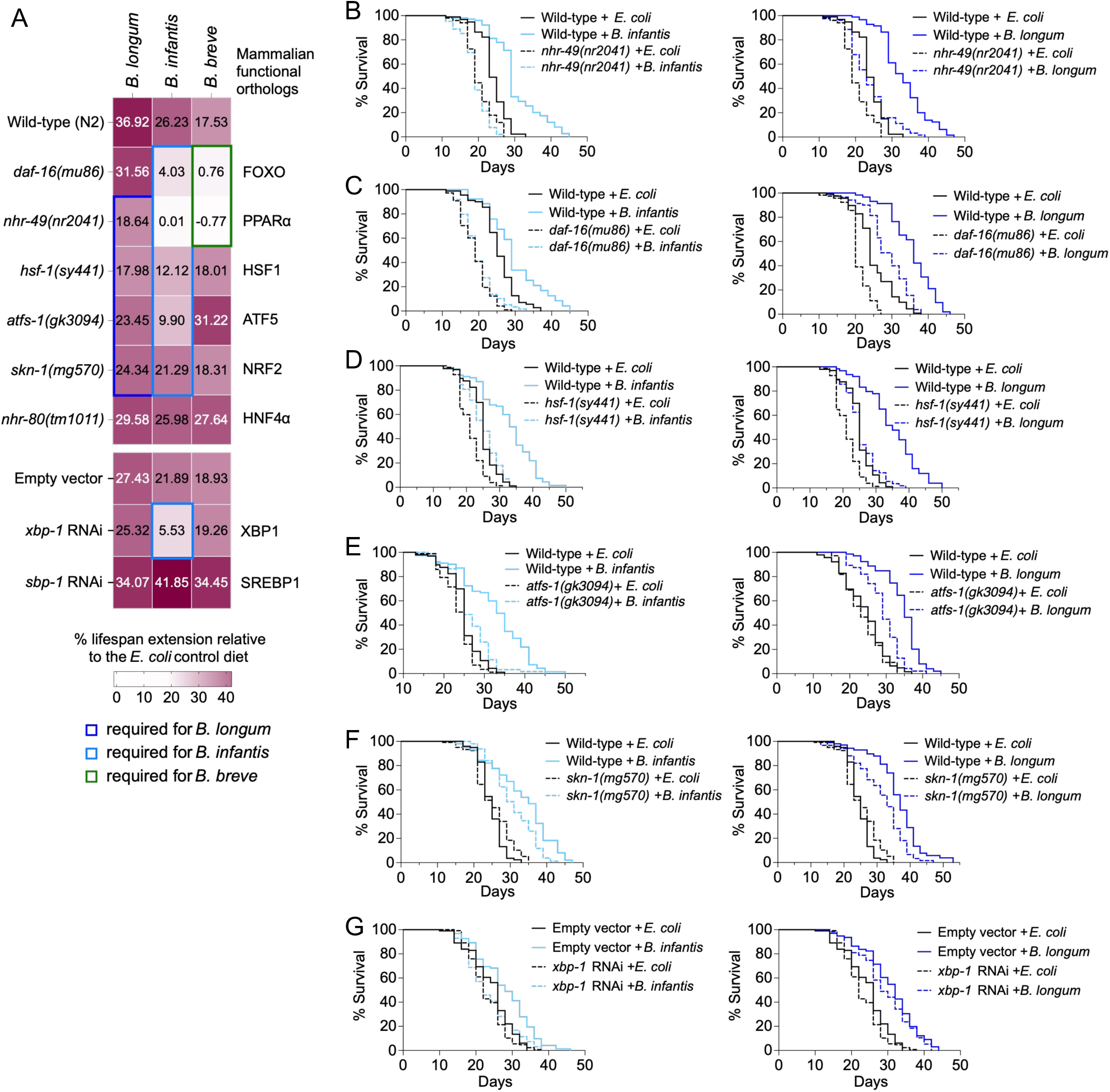
Distinct *Bifidobacteria* strains regulate host lifespan via shared and divergent molecular aging pathways. **A**. Lifespan summary heatmap. Each row indicates % lifespan extension of worms fed different *Bifidobacteria* strains comparing to the *E. coli* control. Each column indicates % lifespan extension in wild-type or transcription factor (TF)-deficient worms, either by mutation (top panel) or RNAi knockdown (bottom panel). Values represent the mean % lifespan extension averaged across two independent experiments, except for *atfs-1(gk3094)* mutants fed on *B. breve*, *nhr-80(tm1011)* fed on *B. longum*, and *sbp-1* RNAi worms fed *B. infantis*, which are based on a single experiment. *B. longum* (dark blue box), *B. infantis* (light blue box), and *B. breve* (green box) require both shared and distinct TFs to achieve full lifespan extension (S3 Table). *xbp-1* RNAi was administered from egg lay to day 1 of adulthood, before transfer to heat-killed diets. Because *sbp-1* RNAi impaired development, it was administered for one day beginning on adult day 1, followed by transfer to heat-killed diets on adult day 2. **B-G**, **left panels**: Lifespan extension by *B. infantis* is reduced in worms deficient in the following TFs, either by mutation or RNAi knockdown: *nhr-49(nr2041)* (-5.46% vs. 25.17% wild-type, n ≥ 92, **B**), *daf-16(mu86)* (3.97% vs. 16.11% wild-type, n ≥ 26, **C**), *hsf-1(sy441)* (17.56% vs. 30.68% wild-type, n ≥ 79, **D**), *atfs-1(gk3094)* (13.03% vs. 30.68% wild-type, n ≥ 78, **E**), *skn-1(mg570)* (21.99% vs. 34.58% wild-type, n ≥ 100, **F**), and *xbp-1* RNAi (2.23% vs. 15.48% empty vector, n ≥ 95, **G**). *P* ≤ 0.0111 for all pairwise interactions, two-way ANOVA. **B-G**, **right panels**: Lifespan extension by *B. longum* is reduced in worms deficient in the following TFs: *nhr-49(nr2041)* (16.53% vs. 37.69% WT, n ≥ 79, **B**), *hsf-1(sy441)* (20.18% vs. 40.38% WT, n ≥ 58, **D**), *atfs-1(gk3094)* (23.11% vs. 42.65% WT, n ≥ 72, **E**), *skn-1(mg570)* (25.02% vs. 48% WT, n ≥ 99, **F**). *n* ≥ 58 per condition. *P* ≤ 0.0011 for all pairwise interactions, two-way ANOVA. The *daf-16(mu86)* mutants (38.47% vs, 40.87% WT, n ≥ 72, **C**) or *xbp-1* RNAi (24.48% vs. 26.93% empty vector, n ≥ 42, **G**) did not significantly affect lifespan extension by *B. longum* (two-way ANOVA). **B-G**, representative of two independent experiments.

All three strains required NHR-49 for their full lifespan effects, although the strength of this requirement differed among strains (Fig 2A and 2B; S1D Fig). Loss of *nhr-49* abolished lifespan extension by *B. infantis* and *B. breve*. By contrast, *nhr-49* loss partially reduced the benefit conferred by *B. longum*, consistent with additional mechanisms contributing to the effect of this strain.

Beyond NHR-49, the strains differed substantially in their pathway requirements. *B. infantis* showed the broadest dependence, requiring NHR-49, DAF-16, HSF-1, ATFS-1, SKN-1, and XBP-1 (Fig 2B–2G, left panels). *B. longum* required NHR-49, HSF-1, ATFS-1, and SKN-1 but retained its lifespan benefit in animals deficient for DAF-16 or XBP-1 (Fig 2B–2G, right panels). *B. breve* required NHR-49 and DAF-16 (S1D and S1E Fig). Thus, the three strains shared dependence on the lipid regulator NHR-49 while requiring different combinations of conserved cytoprotective regulators.

Neither *nhr-80* mutation nor *sbp-1* RNAi significantly reduced lifespan extension by any of the three strains (Fig 2A and S3 Table). Reduced *sbp-1* expression was confirmed following RNAi treatment (S1F Fig). Loss of *hlh-30*, however, did more than eliminate the lifespan benefit. In *hlh-30* mutants, each *Bifidobacteria* diet shortened lifespan below that of the corresponding *E. coli* control (S1G–S1I Fig), revealing a diet-specific vulnerability to *Bifidobacteria* exposure. We therefore focused subsequent mechanistic analyses on regulators whose loss attenuated the bacterial benefit without producing the diet-specific lifespan shortening observed in *hlh-30* mutants.

### The selected *Bifidobacteria* strains differ in mTOR pathway interactions but share mitochondrial stress response interactions

We next examined genetic interactions with established dietary restriction, mTOR, and mitochondrial longevity pathways. We first tested *eat-2* mutants, a genetic model of dietary restriction^37^; *raga-1* and *rsks-1* mutants, which disrupt components of mTORC1 signaling^38,39^; and *rict-1* mutants, which impair mTORC2 signaling^40^. *B. infantis* and *B. breve* further extended lifespan in each of these long-lived genetic backgrounds (S1J–S1M Fig). By contrast, the lifespan benefit conferred by *B. longum* was reduced in *eat-2* mutants (21.98% versus 34.99% in wild type), *raga-1* mutants (16.80% versus 42.65%), and *rsks-1* mutants (31.58% versus 48%). Its lifespan benefit was not reduced in *rict-1* mutants (46.31% versus 34.99% in wild type; *P* < 0.05 for all four comparisons by two-way ANOVA; S1J–S1M Fig). Because *eat-2* longevity is mediated in part by reduced mTORC1 signaling, the combined results suggest that *B. infantis* and *B. breve* act largely independently of the tested mTORC1 and mTORC2 longevity pathways. By contrast, the reduced benefit of *B. longum* in *eat-2*, *raga-1*, and *rsks-1* mutants, but not *rict-1* mutants, indicates partial overlap with mTORC1-associated mechanisms.

Because *B. infantis* and *B. longum* both required ATFS-1 for full lifespan extension, we next examined their effects in the long-lived, mitochondrial electron-transport-chain mutants *nuo-6* and *isp-1*^41^. These mutations impair complex I and complex III activity, respectively, and activate ATFS-1-dependent mitochondrial stress responses^29^. The lifespan benefits of both strains were reduced in these mutant backgrounds compared with wild-type animals. For *B. longum*, lifespan extension was reduced from 35.4% in wild type to 13.44% in *nuo-6* mutants and from 38.62% to 5.28% in *isp-1* mutants. For *B. infantis*, lifespan extension was reduced from 26.08% in wild type to 14.88% in *nuo-6* mutants and 10.03% in *isp-1* mutants (S1N Fig). These results indicate that the effects of both strains partially overlap with mitochondrial stress response pathways.

Collectively, these analyses reveal distinct interactions with mTOR-associated longevity pathways but shared interactions with mitochondrial stress response pathways. *B. infantis* and *B. breve* continued to extend lifespan in the mTORC1 and mTORC2 mutant backgrounds, whereas *B. longum* showed partial dependence on mTORC1-associated longevity mechanisms. *B. infantis* and *B. longum* also showed similar genetic interactions with mitochondrial electron transport chain mutants, consistent with their shared requirement for ATFS-1.

### *B. infantis*-induced lifespan extension and oxidative stress protection require distinct but overlapping host regulators

We next asked whether the host regulators required for *B. infantis*-dependent lifespan extension also mediated protection from oxidative stress. We measured survival after paraquat exposure in animals deficient for *atfs-1*, *hsf-1*, *skn-1*, *nhr-49*, *daf-16*, or *xbp-1*, using the same genetic conditions tested in the lifespan analyses (Fig 3A–3D; S4 Table). Loss of NHR-49 or XBP-1 significantly suppressed the oxidative stress protection conferred by *B. infantis*, whereas loss of ATFS-1 or SKN-1 produced smaller reductions that did not reach statistical significance under these conditions. By contrast, loss of DAF-16 or HSF-1 did not impair paraquat survival on the *B. infantis* diet (S2A and S2B Fig). Thus, the pathway requirements for lifespan extension and oxidative-stress protection overlapped but were not identical. NHR-49, ATFS-1, SKN-1, and XBP-1 were required for both phenotypes, whereas DAF-16 and HSF-1 were specifically required for the lifespan benefit.

**Fig 3.**
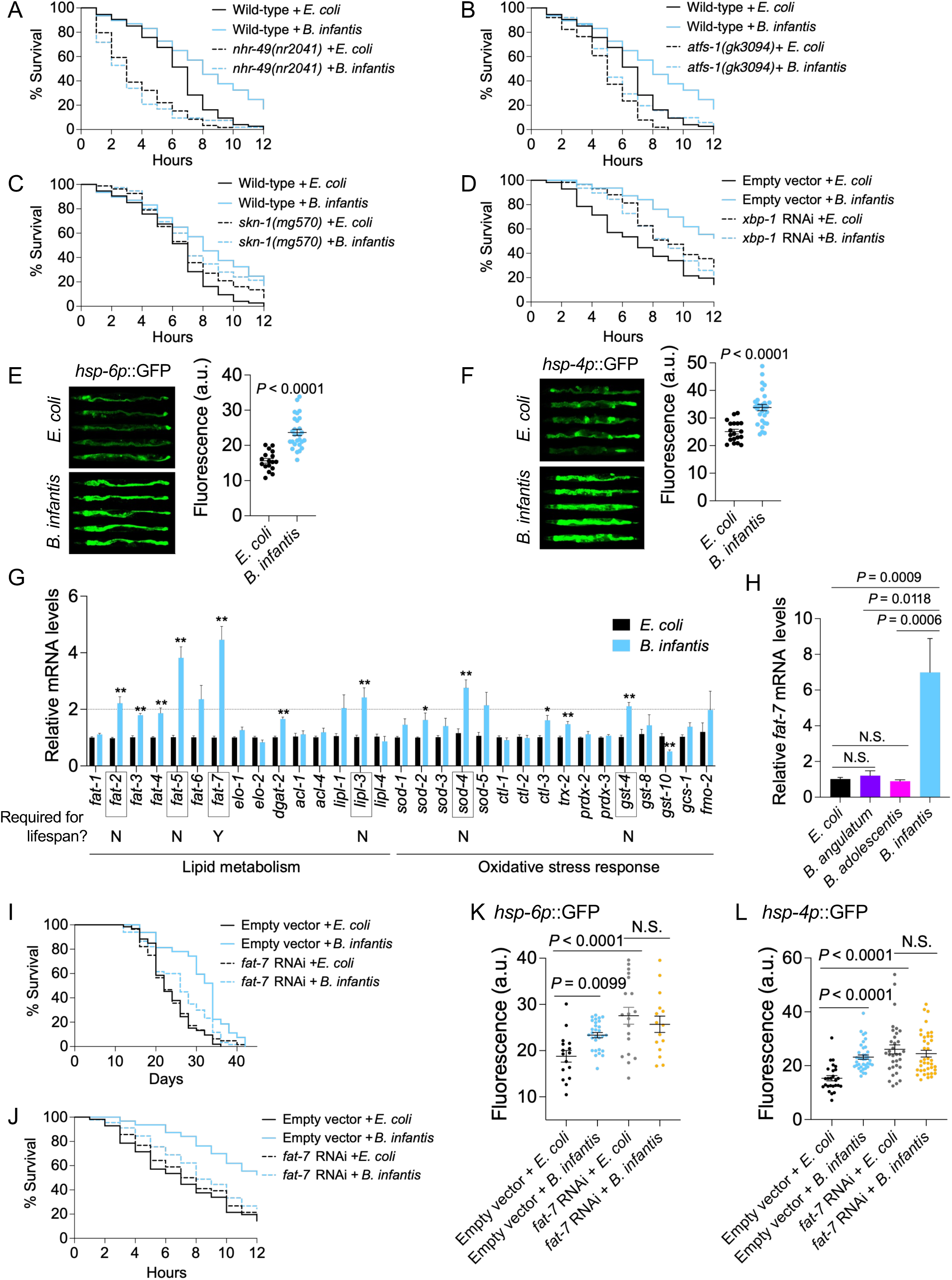
*Bifidobacteria*-responsive TFs coordinate lifespan and stress responses. **A-D.** Enhanced oxidative stress protection conferred by *B. infantis* is reduced in the following mutants relative to wild-type (WT) or in RNAi-treated worms relative to empty vector (EV) controls: *nhr-49(nr2041)* (-7.49% vs. 25.99% WT, *P* = 0.0065), *atfs-1(gk3094)* (18.70% vs. 25.99% WT, N.S.), *skn-1(mg570)* (7.24% vs. 25.99% WT, N.S.), and *xbp-1* RNAi (-4.04% vs. 43.75% EV, *P* < 0.0001). *n* ≥ 51 per condition. *P* values: two-way ANOVA. Representative of two independent experiments. **E-F**. Representative images and quantification of whole worm GFP fluorescence from *hsp-6p*::GFP (**E**) and *hsp-4p*::GFP (**F**) transcriptional reporters in middle-aged (adult day 8) worms. Mean ± s.e.m., *n* ≥ 18 per condition, representative of three independent experiments. *P* values: two-tailed Mann-Whitney. **G.** RT-qPCR analysis of a panel of key enzymes regulating lipid metabolism or oxidative stress response. Mean ± s.e.m from 2-3 independent experiments, each with 3 biological replicates. *P* values: two-tailed Mann-Whitney with Benjamini-Hochberg correction for multiple comparisons (\*\**P* < 0.01, \**P* < 0.05). A threshold of ≥2-fold change (adjusted *P* < 0.05) was used to select candidate enzymes (*fat-2*, *fat-5*, *fat-7*, *lipl-3*, *sod-4*, and *gst-4*) for functional testing of their individual roles in *B. infantis-*mediated lifespan extension. Y indicates significant suppression of lifespan extension by candidate gene mutation or RNAi knockdown; N indicates no significant suppression by two-way ANOVA. *fat-7* RNAi suppressed lifespan extension (*P* < 0.0001), whereas mutation of *fat-5*, *fat-7*, or *sod-4* and RNAi knockdown of *fat-2*, *lipl-3,* or *gst-4* did not significantly suppress lifespan extension. All RNAi treatments were administered from egg lay to day 1 of adulthood before transfer to heat-killed diets, except *fat-2* RNAi. Because *fat-2* RNAi impaired development, it was administered for one day beginning on adult day 1, followed by transfer to heat-killed diets on adult day 2. **H.** RT-qPCR analysis of *fat-7* mRNA levels in worms fed *E. coli*, *B. infantis*, and non-lifespan-extending *Bifidobacteria* strains. *P* values: two-tailed Mann-Whitney with Benjamini-Hochberg correction for multiple comparisons. **I-J**. *fat-7* RNAi significantly reduces *B. infantis-*mediated lifespan extension (9.57% vs. 32.07% EV controls; n ≥ 32, *P* = 0.0059, **I**) and oxidative stress protection (9.73% vs. 43.75% EV controls; n ≥ 45, *P* = 0.0132, **J**). *P* values: two-way ANOVA. **K-L**. Quantification of whole worm GFP fluorescence of *hsp-6p*::GFP (**K**) and *hsp-4p*::GFP (**L**) transcriptional reporters in middle-aged (adult day 8) worms treated with empty vector (EV) control or *fat-7* RNAi. Mean ± s.e.m, n ≥ 16 worms per condition. *P* values: Kruskal-Wallis with Benjamini-Hochberg correction for multiple comparisons. Representative of two independent experiments.

We next examined whether *B. infantis* activated transcriptional programs associated with these pathways. Expression of transcriptional reporters for the ATFS-1 target *hsp-6*^42^ and the XBP-1 target *hsp-4*^43^ increased in animals fed *B. infantis* relative to *E. coli* controls (Fig 3E and 3F). We also measured a targeted panel of genes involved in fatty acid metabolism, complex lipid synthesis, lipid breakdown, and cellular stress responses (Fig 3G). In young adults, *B. infantis* increased the expression of several lipid metabolism genes by at least twofold, including the delta-9 desaturases *fat-5* and *fat-7*, the polyunsaturated fatty acid desaturase *fat-2*, and the lysosomal lipase *lipl-3*. The antioxidant genes *sod-4* and *gst-4* were also induced.

Because *fat-7* encodes an NHR-49-regulated delta-9 desaturase, we examined whether its induction distinguished *B. infantis* from strains that did not extend lifespan. *B. infantis* strongly increased *fat-7* expression, whereas *B. adolescentis* and *B. angulatum* did not significantly increase *fat-7* expression relative to the *E. coli* control (Fig 3H). This strain-level association implicated FAT-7-dependent fatty acid desaturation as a candidate mechanism linking bacterial exposure to host lipid metabolism and physiology. Increased delta-9 desaturase expression can accompany triglyceride accumulation under some physiological conditions^44,45^. We therefore measured neutral lipid abundance using Oil Red O staining^46^. Animals fed *B. infantis* showed reduced rather than increased staining relative to *E. coli* controls (S2C Fig). Thus, induction of lipid metabolism genes was not accompanied by increased neutral lipid storage but instead occurred alongside reduced neutral lipid abundance, suggesting altered host lipid metabolism.

This distinction is important because lipid composition, rather than bulk neutral lipid accumulation, can influence mitochondrial membrane properties^47^, susceptibility to lipid peroxidation^48^, oxidative stress resistance^49,50^, and longevity^34,44^.

### *B. infantis* and *B. longum* require FAT-7 for full lifespan extension and oxidative stress protection

To determine whether genes induced at least twofold by *B. infantis* were functionally required for lifespan extension, we performed a targeted genetic screen using available mutants and RNAi against selected lipid metabolism genes (*fat-5*, *fat-7*, *fat-2*, and *lipl-3*) and antioxidant genes (*sod-4* and *gst-4*). Individual mutations in *fat-5*, *fat-7*, or *sod-4*, or RNAi targeting *fat-2*, *lipl-3*, or *gst-4*, did not significantly suppress *B. infantis*-dependent lifespan extension (S3 Table). However, FAT-5, FAT-6, and FAT-7 have overlapping roles in monounsaturated fatty acid (MUFA) synthesis^51^. FAT-5 preferentially produces C16:1, whereas FAT-6 and FAT-7 primarily produce C18:1. Consistent with this functional redundancy, combined loss of *fat-6* and *fat-7* significantly attenuated *B. infantis*-dependent lifespan extension (S2D Fig).

Additionally, *fat-7* RNAi reduced both the lifespan extension and oxidative stress protection conferred by *B. infantis* (Fig 3I and 3J). The stronger RNAi phenotype may reflect a more acute reduction in desaturase function before compensatory responses by related desaturases can develop, as may occur in the stable *fat-7* mutant. By contrast, *fat-2* RNAi did not significantly reduce lifespan extension (S3 Table), indicating that FAT-2-dependent conversion of C18:1 to C18:2 was not required under these conditions. This finding does not exclude contributions from downstream polyunsaturated fatty acid (PUFA) metabolism, including C20 PUFAs synthesized from dietary C18:2 pools. Together, these results identify FAT-7-dependent delta-9 fatty acid desaturation as an important contributor to *B. infantis*-dependent longevity and oxidative stress protection.

We next asked whether this mechanism was shared by *B. longum*, which differed from *B. infantis* in several upstream pathway requirements. Like *B. infantis*, *B. longum* increased *fat-5* and *fat-7* expression, although the magnitude of induction was smaller (S2E Fig). Deficiency in *nhr-49*, *hsf-1*, *atfs-1*, and *skn-1* differentially altered *B. longum*-dependent *fat-7* expression (S2F Fig), indicating that multiple regulators required for lifespan extension also modulate desaturase expression^52–54^. Individual *fat-5* or *fat-7* mutations did not significantly reduce *B. longum*- dependent lifespan extension (S3 Table). In contrast, the *fat-6*;*fat-7* double mutation and *fat-7* RNAi both attenuated the lifespan benefit (S2G and S2H Fig). Additionally, *fat-7* RNAi suppressed the oxidative stress protection conferred by *B. longum* (S2I Fig). Thus, despite differing in their upstream cytoprotective pathway requirements, *B. infantis* and *B. longum* both require FAT-7-dependent delta-9 fatty acid desaturation for full lifespan extension and oxidative stress protection.

FAT-7 regulates MUFA synthesis and thereby shapes membrane lipid composition, which is important for mitochondrial and endoplasmic reticulum (ER) function and their associated stress responses^55^. Because ATFS-1 and XBP-1 were both required for *B. infantis*-dependent lifespan extension, we examined how loss of FAT-7 affected activation of these cytoprotective pathways by *B. infantis*. The *hsp-6p*::GFP^42^ and *hsp-4p*::GFP^43^ reporters provide readouts of ATFS-1 and XBP-1 pathway activity, respectively. In control animals, *B. infantis* increased expression of both reporters, whereas no further induction was observed following *fat-7* RNAi (Fig 3K and 3L). Notably, *fat-7* RNAi elevated basal expression of both reporters independently of bacterial diet. These findings suggest that loss of FAT-7 activates basal mitochondrial and ER stress responses while limiting their additional induction by *B. infantis*. Thus, FAT-7 may help maintain lipid homeostasis required for the cytoprotective response to the bacterial diet.

Together, these findings establish FAT-7 as a shared functional requirement for the full lifespan extension and oxidative stress protection conferred by *B. infantis* and *B. longum*. They further connect FAT-7-dependent delta-9 fatty acid desaturation to the mitochondrial and ER stress responses induced by *B. infantis*.

### *B. infantis* and *B. longum* induce overlapping FAT-7-dependent lipid signatures

We hypothesized that FAT-7-dependent desaturation contributes to the physiological effects of *Bifidobacteria* diets by remodeling the fatty acyl compositions of complex lipids. Complex lipids influence membrane organization, fluidity, signaling, and organelle function. Although *fat- 7* overexpression and dietary MUFA supplementation can extend *C. elegans* lifespan^44,45^, the complex lipid species associated with FAT-7-dependent lifespan extension in response to beneficial bacterial diets remain poorly defined.

To identify lipid changes associated with lifespan-extending *Bifidobacteria* strains, we performed two untargeted LC-MS/MS lipidomic experiments using established methods^56–58^ optimized on our in-house mass spectrometry platform^59^. Across both experiments, we detected more than 450 host lipid species representing 15 lipid subclasses (S3A Fig). In the first experiment, we compared middle-aged animals fed the three strains that produced the largest lifespan benefits, *B. infantis*, *B. longum*, and *B. breve*, with animals fed the non-lifespan-extending strains *B. adolescentis* and *B. angulatum* (S1 Data). Principal component analysis separated animals fed the lifespan-extending strains, particularly *B. infantis* and *B. longum*, from those fed the two non-lifespan-extending strains, which clustered together (Fig 4A). Similar patterns were observed when the lipidomic data were normalized to either *B. adolescentis* or *B. angulatum* (Fig 4A and S3B Fig). In the second experiment, animals were exposed from hatching to either empty-vector control RNAi or *fat-7* RNAi. On the first day of reproductive adulthood, animals in each RNAi condition were transferred to diets of *E. coli*, *B. infantis*, or *B. longum*, and were collected for lipidomic profiling in middle age (S2 Data). Principal component analysis separated samples according to both bacterial diet and RNAi treatment, indicating that bacterial exposure and FAT-7 activity each contributed to variation in host lipid composition (Fig 4B).

**Fig 4.**
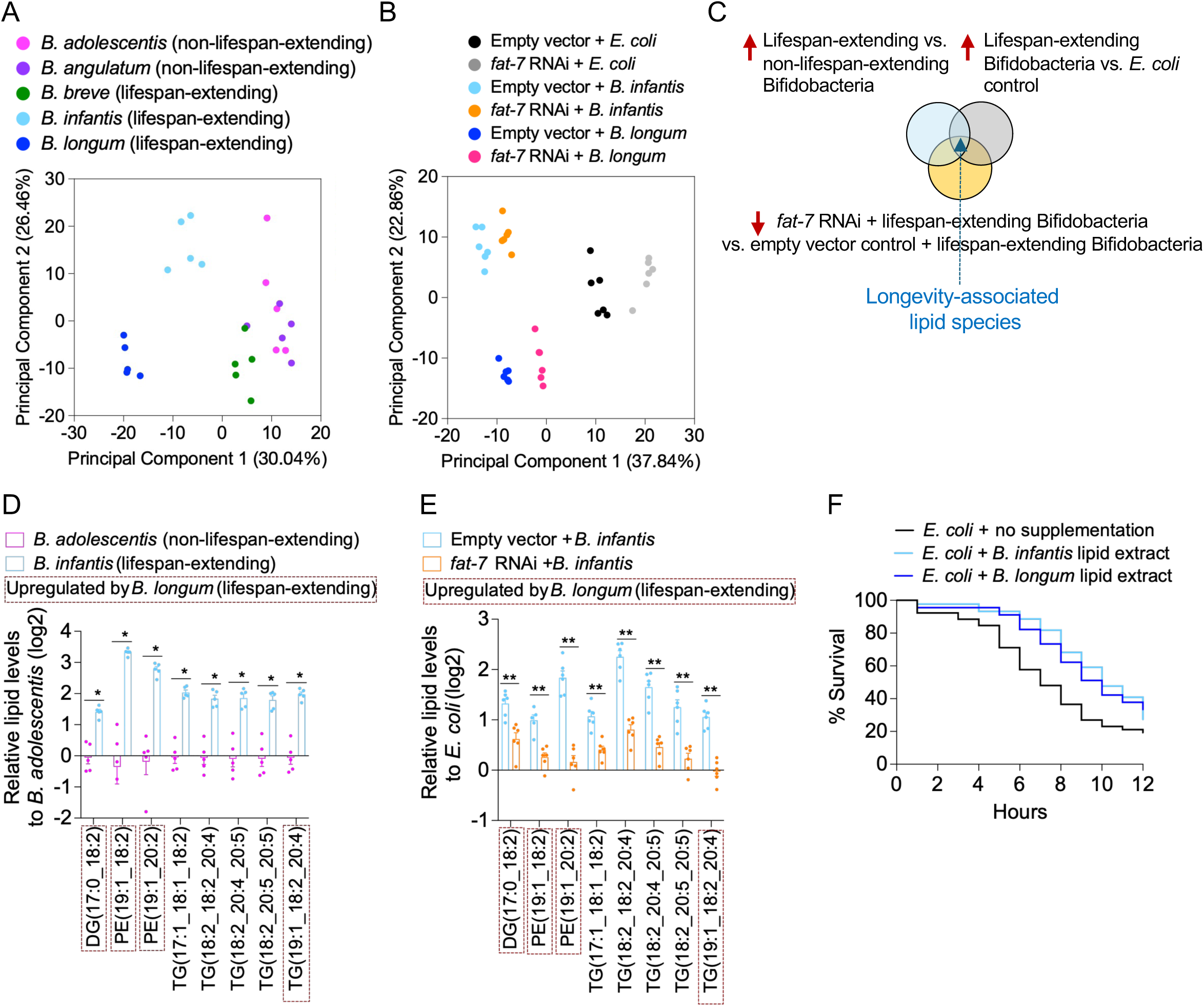
Identification of *Bifidobacteria*-dependent, longevity-associated lipid signatures. Principal component analysis (PCA) separates lipidomic profiles of middle-aged worms based on distinct *Bifidobacteria* diets. Data are log2 fold-change values normalized to the non-lifespan-extending *B. adolescentis* control. **B.** PCA separates lipidomic profiles of middle-aged (adult day 8) worms by bacterial diets and RNAi treatments. Data are log2 fold-change values normalized to the *E. coli* empty vector control. **C**. Schematic illustrating the three filtering criteria used to identify lipid signatures that are both *Bifidobacteria*-dependent and longevity-associated. **D-E**. Lipid species passing all three filtering criteria are shown. These lipids are 1) significantly upregulated by *B. infantis* relative to *B. adolescentis* control (≥2-fold, or log2 ≥1, adjusted *P* < 0.05) (**D**), 2) significantly upregulated by *B. infantis* relative to *E. coli* control under the same threshold (**E**), and 3) significantly reduced by *fat-7* RNAi (adjusted *P* < 0.05) (**E**). Mean ± s.e.m. from one experiment with 5-6 biological replicates per condition, each containing ∼500 worms. Lipid species highlighted in red dashed boxes indicate the overlapping lipids associated with lifespan extension by *B. longum*. *P* values: two-tailed Mann-Whitney with Benjamini-Hochberg correction for multiple comparisons. **F**. Supplementation of bulk lipid extracts from *B. infantis* or *longum* on top of *E. coli* enhances oxidative stress resistance relative to the un-supplemented control. Lipid extracts were resuspended in 50% methanol, dried onto agar plates, and seeded with *E. coli* prior to use; the non-supplemented control contained 50% methanol without lipid extract. n ≥ 44 per condition. *P* ≤ 0.013 for both comparisons by log-rank. Representative of two independent experiments.

We integrated the two datasets to identify candidate lipid species associated with lifespan-extending *Bifidobacteria* diets and FAT-7. Lipids were required to meet three criteria: enrichment in animals fed strains that extended lifespan relative to strains that did not; enrichment relative to *E. coli*-fed controls; and reduction following *fat-7* RNAi (Fig 4C). Enrichment was defined as at least a twofold increase (log₂ fold change ≥ 1) with corrected *P* < 0.05. A focused subset of phosphatidylethanolamine (PE), diacylglycerol (DG), and triacylglycerol (TG) species met all three criteria. These lipids were enriched in *B. infantis*-fed animals relative to both non-lifespan-extending *Bifidobacteria* strains and *E. coli* controls, and their abundance was reduced by *fat-7* RNAi (Fig 4D and 4E; S3C and S3D Fig). Most contained at least one MUFA chain, including C17:1, C18:1, or C19:1, and several also contained PUFA chains, including C18:2, C20:2, C20:4, or C20:5. Complex lipids containing C17:1, C18:1, and C19:1 have previously been detected in *C. elegans* lipidomes^45,60^, supporting the plausibility of these lipid assignments. Although several candidate species contained PUFA chains, *fat-2* RNAi did not reduce the lifespan extension conferred by *B. infantis* (S3 Table). This result suggests that the lifespan phenotype does not depend on host FAT-2-mediated synthesis of C18:2 under these conditions. The PUFA-containing species may instead reflect the incorporation of dietary fatty acids together with downstream host elongation, desaturation, and lipid remodeling.

Applying the same analysis to *B. longum* identified a partially overlapping set of lipid species (S3E–S3H Fig). Lipids shared between *B. infantis-* and *B. longum*-fed animals included PE(19:1_18:2), PE(19:1_20:2), DG(17:0_18:2), and TG(19:1_18:2_20:4) (Fig 4D and 4E; S3C–S3H Fig). PE ether lipids, including PE O- and PE P-type species, have been linked to lifespan extension in *C. elegans*^61^, and increased PE abundance promotes longevity across species by enhancing autophagic flux^62^. By contrast, both shared PE species identified here were non-ether lipids that paired a C19:1 MUFA chain with a PUFA chain. These species therefore define a distinct lipid signature associated with lifespan-extending *Bifidobacteria* strains. These findings identify a shared lipid signature associated with two *Bifidobacteria* strains that differ in several upstream pathway requirements yet share dependence on FAT-7.

We next asked whether lipids isolated from these bacterial strains could recapitulate part of the protective phenotype. Bulk lipid extracts from *B. infantis* or *B. longum* were added to a standard heat-killed *E. coli* diet beginning on the first day of reproductive adulthood. Extracts from either strain increased survival during paraquat exposure relative to the unsupplemented *E. coli* control (Fig 4F), indicating that bacterial lipids can contribute to oxidative stress protection. Collectively, these findings show that *B. infantis* and *B. longum* induce distinct but overlapping changes in host lipid composition that depend in part on FAT-7. They further indicate that host lipid remodeling and bacterially derived lipids make complementary contributions to the protective effects of beneficial *Bifidobacteria* diets against oxidative stress.

## Discussion

Although individual *Bifidobacteria* strains have been reported to extend lifespan or improve stress resistance in *C. elegans*^10,11,63^, whether diverse human-associated strains engage shared or distinct host mechanisms remains unclear. Using an anaerobic bacteria–*C. elegans* platform, we compared 11 strains representing nine human gut *Bifidobacteria* species. *B. infantis* ATCC 15697, *B. longum* NCC 2705, and *B. breve* DSMZ 20213 produced the largest lifespan benefits and improved overlapping measures of physiological resilience, whereas responses to the other strains ranged from smaller benefits to neutral or detrimental effects. These findings establish that host benefits vary substantially among *Bifidobacteria* strains rather than representing a uniform property of the genus. Because animals were transferred as adults to heat-killed, biomass-normalized diets, the observed effects primarily reflect adult host responses to bacterial products rather than live bacterial growth or developmental exposure. Although biomass normalization cannot ensure nutritional equivalence, preserved pharyngeal pumping, strain-specific stress phenotypes, sustained lifespan extension under mixed dietary conditions, and the protective activity of bacterial lipid extracts collectively argue against a simple dietary restriction explanation.

Genetic analyses revealed a shared requirement for NHR-49 among the three strains, linking their lifespan benefits to host lipid regulation. Loss of *nhr-49* abolished lifespan extension by *B. infantis* and *B. breve* and partially reduced the benefit conferred by *B. longum*. Beyond this shared genetic requirement, the strains depended on different combinations of conserved cytoprotective regulators. *B. infantis* showed the broadest dependence, requiring DAF-16, HSF-1, ATFS-1, SKN-1, and XBP-1 in addition to NHR-49. *B. longum* required HSF- 1, ATFS-1, and SKN-1 but not DAF-16 or XBP-1, whereas *B. breve* required DAF-16 together with NHR-49. Building on prior evidence that gut microbes and microbe-derived lipids can alter host lipid metabolism^64–66^, these findings identify NHR-49-dependent host lipid regulation as a shared mechanistic requirement for the lifespan benefits of the three selected *Bifidobacteria* strains.

These pathway requirements also differed across physiological outcomes. For *B. infantis*, NHR-49, ATFS-1, SKN-1, and XBP-1 mediated both lifespan extension and oxidative stress protection, whereas DAF-16 and HSF-1 were required only for lifespan extension. Enhanced oxidative stress resistance may therefore contribute, at least in part, to the lifespan benefit conferred by this strain. Additional genetic interactions further distinguished the strains. *B. longum* showed partial overlap with genetic backgrounds associated with dietary restriction and reduced mTORC1 signaling, whereas *B. infantis* and *B. breve* extended lifespan largely independently of these backgrounds. By contrast, *B. infantis* and *B. longum* showed similar interactions with mitochondrial electron-transport-chain mutants.

Despite these differences, *B. infantis* and *B. longum* both required FAT-7-dependent delta- 9 fatty acid desaturation for lifespan extension and oxidative stress protection. Both strains induced *fat-7* expression, whereas two strains that did not extend lifespan failed to do so.

Moreover, *fat-7* RNAi attenuated both phenotypes in animals fed either strain. Previous studies have shown that FAT-7 and MUFAs influence *C. elegans* longevity^44,45,51,52^ and that bacterial lipid signals can regulate NHR-49-dependent *fat-7* expression^67^. Our findings extend these findings by identifying FAT-7-dependent delta-9 fatty acid desaturation as a shared functional requirement for the full benefits of two lifespan-extending *Bifidobacteria* strains with otherwise different host pathway dependencies.

Untargeted lipidomics further linked this shared genetic requirement to host lipid composition. Lifespan-extending strains induced lipid profiles distinct from those produced by strains that did not extend lifespan or by the *E. coli* control. A focused set of PE, DG, and TG species was enriched by *B. infantis* or *B. longum* and reduced by *fat-7* RNAi. The signatures induced by the two strains were distinct but partially overlapping, consistent with their different upstream requirements and shared dependence on FAT-7. Bulk lipid extracts from either strain also increased oxidative stress resistance when added to a standard *E. coli* diet, supporting a functional contribution from bacterial lipids to oxidative stress protection. Together, these findings provide a foundation for identifying the active bacterial lipid species and defining how they interact with host lipid remodeling. Future supplementation studies with individual lipids and isotope tracing could link specific microbial lipids to the host lipid signatures and physiological benefits.

Together, our findings support a strain-selective model of partial mechanistic convergence. Related human gut *Bifidobacteria* strains differed in their physiological effects and cytoprotective pathway requirements, yet the three strongest lifespan-extending strains all depended on NHR-49 to varying degrees. *B. infantis* and *B. longum* also required FAT-7 for lifespan extension and oxidative stress protection and induced partially overlapping host complex lipid signatures. This work identifies gut bacteria–host lipid interactions as a mechanistic axis linking microbial products, stress resilience, and longevity. Given the conservation of these host lipid metabolism pathways, comparable mechanisms may contribute to the effects of *Bifidobacteria* on mammalian healthspan. These findings provide a foundation for testing which strains and bacterial molecules act through these pathways in more complex hosts.

## Materials and Methods

### Worm strains

All *Caenorhabditis* elegans strains were maintained at 20°C on standard nematode growth medium (NGM) agar plates seeded with *Escherichia coli* (*E. coli*) OP50, unless otherwise indicated. All worm strains were obtained from the *Caenorhabditis* Genetics Center (CGC). A complete list of strains used in this study is provided in S5 Table. All experiments were performed with N2 hermaphrodite worms.

### Bacterial culture

The bacterial strains used in this study are listed in S1 Table. All *Bifidobacteria* strains were cultured in De Man-Rogosa-Sharpe (MRS) medium at 37°C under anaerobic conditions in a Coy Laboratories chamber maintained at an atmosphere of approximately 80% N₂, 15% CO₂, and 5% H₂. To synchronize bacterial growth, individual colonies of each strain was inoculated into 1.5 mL of MRS medium in five replicates using 2 mL 96-well blocks and incubated overnight (16- 18 hours) anaerobically at 37°C. These pre-cultures were sub-cultured into fresh MRS medium at a 1:50 dilution (800 µL into 40 mL) in individual 50 mL Falcon tubes and incubated anaerobically for another 16-18 hours at 37°C to reach early saturation. In parallel, 200 µL of each subculture was grown in a microplate reader (BioTek Epoch 2) to monitor cell growth by measuring optical density at 600 nm. *E. coli* OP50 was cultured under aerobic conditions in Luria-Bertani (LB) broth at 37°C with agitation (200 rpm) for 18 hours. For all *Bifidobacteria* and *E. coli* cultures, cells were harvested at early stationary phase and heat-killed in a 65°C water bath for 30 minutes. Cell pellets were collected by centrifugation (4,500 × g) for 10 minutes, washed twice with sterile M9 buffer (22 mM KH_2_PO_4_, 42 mM Na_2_HPO_4_, 86 mM NaCl and 1mM MgSO_4_, dissolved in water), and resuspended in M9 to normalize bacterial biomass across strains to 0.2 mg wet weight per uL. The concentrated bacterial suspensions were aliquoted and stored at -80°C until use.

### RNA interference

RNA interference (RNAi) was performed as previously described^44^ by feeding worms *E. coli* HT115 expressing double-stranded RNA (dsRNA) targeting *fat-2*, *fat-7*, *sbp-1*, *xbp-1*, *lipl-3*, and *gst-4* (Ahringer or Vidal RNAi libraries). All constructs were sequence-verified prior to use.

Bacteria carrying the empty vector L4440 served as controls. Overnight cultures grown in LB broth with ampicillin (50 μg/mL) at 37°C were induced with isopropylthiogalactoside (IPTG; 0.4 mM) for 2 h, concentrated 40-fold, and stored at 4°C for up to two weeks. Concentrated bacteria were seeded onto NGM plates supplemented with ampicillin (50 μg/mL) and IPTG (0.4 mM).

Unless otherwise noted, RNAi treatment began at hatching and continued until the first day of adulthood before switching onto heat-killed diets. Because *fat-2* and *sbp-1* RNAi impaired development, they were administered for one day beginning on adult day 1, followed by transfer to heat-killed diets on adult day 2.

### Lifespan assay

Worm populations were synchronized by placing young adult worms to lay eggs on NGM plates seeded with live *E. coli* OP50 for 4-6 hours, after which the adults were removed. All worms were grown on live *E. coli* until day 1 of adulthood (typically within 72 hours post-egg laying) to ensure synchronized development across all conditions. Upon reaching adulthood, worms were transferred to NGM plates containing ampicillin (50 µg/mL) and 5-fluoro-2′-deoxyuridine (FUdR; 50 µM) seeded with either heat-killed *E. coli* OP50 or heat-killed *Bifidobacteria* strains.

Antibiotics were included to eliminate residual live bacterial carryover onto heat-killed diets, and FUdR was included to reduce censoring due to internal hatching (“bagging”). Worms were transferred onto freshly seeded plates every other day and scored as alive or dead based on movement in response to gentle prodding with a worm pick. Individuals that crawled off the plate or died from vulval rupture or bagging were censored from analysis. For all lifespan plots, the day of hatching was designated as day 1, except for experiments using the mitochondrial *isp- 1* and *nuo-6* mutants, where day 1 corresponded to the first day of adulthood due to their extended developmental period relative to wild-type worms. Kaplan-Meier survival curves were generated using GraphPad Prism, and statistical significance was assessed by log-rank tests in JMP software. Two-way ANOVA was used to determine lifespan interactions between experimental conditions. Representative Kaplan-Meier curves are shown in the main and supporting information, and summary of lifespan statistics is provided in S3 Table.

### Pharyngeal pumping and locomotion activity measurement

Synchronized adult day 1 worms were transferred to plates seeded with either heat-killed *Bifidobacteria* strains or *E. coli* control and maintained by transferring to freshly seeded plates every other day until the day of assay. Pharyngeal pumping rates were measured by counting the number of contractions of the terminal pharyngeal bulb over a 30-second interval in worms actively feeding within the bacterial lawn. Measurements were performed using a dissecting microscope, with a minimum of 10 worms analyzed per biological replicate. For locomotor activity measurements, a single worm was placed into a drop of M9 buffer on an NGM plate without bacteria, and the number of body bends (sine waves) completed within 30 seconds was recorded. Two plates were used per condition, with at least 30 worms per plate. Statistical significance was determined using the Kruskal-Wallis test followed by Benjamini–Hochberg correction for multiple comparisons.

### Heat, cold, and oxidative stress assays

Worms were age-synchronized and maintained as described previously. Worms were fed on plates seeded with either heat-killed *Bifidobacteria* strains or *E. coli* control until adult day 8 of adulthood (middle-age), when they were subjected to the indicated stress conditions. During all stress assays, worms were maintained on the same bacterial diet provided prior to stress exposure until the end of the experiment. For heat stress, worms were shifted from 20°C (maintenance) to 34°C and remained at that temperature for the entire duration of this experiment. Survival was recorded every 4 hours until all worms in the control group had died. Two plates per condition were used, with at least 30 worms per plate. Kaplan-Meier survival curves were generated using GraphPad Prism, and statistical significance was assessed by log-rank tests in JMP software. For cold stress, worms were shifted from 20°C to 4°C for 60 hours, then returned to 20°C for a 24- hour recovery period before survival was scored as the percentage of total worms per condition. Eleven replicates across two independent experiments were performed, with 15-25 worms per plate. Statistical significance was determined using the Kruskal-Wallis test followed by Benjamini-Hochberg correction for multiple comparisons. For oxidative stress, worms were transferred to NGM plates containing 200 mM paraquat and kept on the same plates until the end of the experiment. Survival was scored hourly for up to 12 hours, by which time most or nearly all wild-type worms fed the *E. coli* control diet had died. Worms that were still alive at the end of experiment were censored. Paraquat plates were prepared one day before use and stored in the dark to minimize oxidation. Two plates were used per condition, with at least 20 worms per plate. Kaplan-Meier survival curves were generated using GraphPad Prism, and statistical significance was assessed by log-rank tests in JMP software. Two-way ANOVA was used to determine the significance of interactions between experimental conditions. Representative Kaplan-Meier curves are shown in the main and supporting information, and summary of oxidative stress survival statistics are provided in S4 Table.

### ROS quantification

Age-synchronized *C. elegans* were maintained at 20 °C and transferred at adult day 1 onto NGM plates seeded with either heat-killed *Bifidobacteria* strains or *E. coli* OP50 (control). Worms were maintained on these diets until adult day 8, at which point they were harvested and washed three times with M9 buffer to remove residual bacteria. Worms were then lysed by sonication on ice in M9 buffer using a QSonica Ultrasonic Processor, and lysates were clarified by centrifugation at 10,000 × *g* for 5 min. Supernatants were collected and protein concentrations was determined using the Pierce BCA Protein Assay Kit (Thermo Fisher Scientific); all samples were normalized to equal protein content prior to ROS measurements. General oxidative activity in worm lysates was measured using the fluorescent probe 2’,7’-dichlorodihydrofluorescein diacetate (H₂DCFDA). Normalized lysates were incubated with H₂DCFDA at a final concentration of 25 µM in black 96-well plates, and fluorescence was measured using a SpectraMax M5*^e^*microplate reader. Mitochondrial superoxide levels were assessed using MitoSOX Red (Thermo Fisher Scientific). Live worms were incubated with MitoSOX Red for 1 hour, washed three times with M9 buffer, and immobilized with 5 mM levamisole on 2% agarose pads. Images were acquired using an Andor Dragonfly spinning-disk confocal microscope, with acquisition focused on the head region. Fluorescence intensity was quantified in ImageJ by manually drawing a region of interest (ROI) around the terminal bulb of the pharynx, a mitochondria-rich tissue. Statistical significance was assessed using the Kruskal-Wallis test followed by Benjamini–Hochberg correction for multiple comparisons.

### GFP imaging and quantification

The expression of *hsp-4* and *hsp-6* was measured on adult day 8 using the transgenic reporter strains SJ4005 [zcls4(*hsp4p*::GFP)] and SJ4100 [zcls13(*hsp-6p*::GFP)], respectively. For fluorescence reporter measurements, worms were induced for 4 h in M9 buffer containing 50 ng/mL tunicamycin. Worms were then anesthetized in M9 buffer containing 500 µM levamisole and mounted on 2% agar pads. Images were acquired using a Leica M165 FC stereomicroscope equipped with a Leica DMC6200 Digital Camera and LAS X Life Science Microscope Software, with identical exposure settings applied across all conditions within each experiment. Image quantification was performed using FIJI. Raw images were background-subtracted and thresholded to define worm boundaries, with consistent threshold values applied across all conditions. Mean GFP fluorescence intensity of each worm, expressed in arbitrary units (a.u.), was measured using the Analyze Particles function and plotted in GraphPad Prism. Statistical significance was assessed using the Kruskal-Wallis test followed by Benjamini–Hochberg correction for multiple comparisons.

### Oil-Red-O staining and quantification

Oil-Red-O (ORO) staining of fixed worms was performed as previously described^44^. Stained worms were mounted on 2% agar pads and imaged at 10× magnification using the same Leica M165 FC stereomicroscope equipped with a Leica DMC6200 Digital Camera and LAS X Life Science Microscope Software, as described above. Images were saved as TIFF files, and identical exposure settings were applied across all conditions within each experiment. Image quantification was performed using FIJI. Raw images were background-subtracted, converted to grayscale, inverted, and thresholded to define worm boundaries, using consistent threshold values across all conditions. Mean ORO intensity per worm (a.u.) was measured using the Analyze Particles function, plotted in GraphPad Prism, and statistical significance was determined using a Kruskal-Wallis test followed by Benjamini–Hochberg correction for multiple comparisons.

### Quantitative RT-PCR

At least 50 age-synchronized worms per condition were collected in biological triplicates at adult day 2 following 24 hours of feeding on heat-killed *Bifidobacteria* strains vs. *E. coli* controls.

Total RNA was extracted and quantified as previously described^44^, treated with DNase I (Amp Grade, New England Biolabs), and reverse-transcribed using Oligo(dT) primers and AccuScript High-Fidelity reverse transcriptase (Agilent) according to the manufacturer’s instructions. RT-qPCR was performed on diluted cDNA using a BioRad thermal cycler and Brilliant III SYBR Green Master Mix (Agilent) or PowerUp SYBR Green Master Mix (Applied Biosystems).

Primers were designed to span exon–exon junctions near the 3′ end of each gene and used at final concentrations of 200 nM (Brilliant III) or 500 nM (PowerUp), following the manufacturers’ instructions. All RT-qPCR primer sequences used in this study are listed in S6 Table. Relative gene expression levels were calculated using the standard ΔΔCₜ method. For each biological replicate, the mean Cₜ value of three technical replicates was used. *act-1* served as the internal reference gene for all analyses.

### Untargeted lipidomics by LC-MS/MS

Lipid extraction^68^ and LC-MS/MS methods^56–58^ were adapted from previously published workflows and optimized on our in-house mass-spectrometry platform. Individual steps on worm lipid extraction, liquid chromatography, mass spectrometry, data-dependent MS2 data acquisition, and data analysis are described in detail below.

### Worm lipid extraction for lipidomics

Methanol (MeOH) and methyl tert-butyl ether (MTBE) were pre-chilled at −20°C and water at 4°C prior to use. Flash-frozen worm pellets stored in 200 µL of M9 buffer were thawed on ice and mechanically lysed using a motorized mortar and pestle tissue grinder for 30 s, with tubes maintained on ice throughout to minimize heat-induced degradation. Next, 60 µL of each lysate was transferred to a 2 mL screwcap vial. Lipids were extracted by sequential addition of 1 µL of equiSPLASH deuterium-labeled lipid standard mix (Avanti Polar Lipids), 224 µL of ice-cold MeOH, and 750 µL of ice-cold MTBE, followed by vortexing for 6 min at 4°C on a Microplate Genie vortexer. Phase separation was initiated by addition of 187.5 µL of water and brief vortexing for 20 s, after which samples were centrifuged at 14,000 × *g* for 2 min at room temperature. Approximately 750 µL of the upper organic phase was carefully transferred to individual amber glass vials while avoiding the interphase layer.

Extracts were dried under a nitrogen stream at 45 L/min in a TurboVap96 evaporator (Biotage) for 15 min at room temperature in the dark. Dried lipid films were resuspended in 75 µL of LC-MS grade MeOH by vortexing for 30 s, then transferred to 150 µL glass microinserts placed inside 2mL amber glass vials. Samples were stored at −20°C until LC-MS/MS analysis.

### Liquid chromatography

Reverse-phase chromatographic separation was performed on a Vanquish Horizon UHPLC system (Thermo Fisher Scientific) equipped with an Accucore C18 column (2.1 × 150 mm, 2.6 μm particle size; Thermo Fisher Scientific). The column temperature was maintained at 45 °C, the autosampler temperature at 15 °C, and the injection volume was 5 μL. The operating pressure ranged from 200 to 350 bar. Mobile phase A consisted of acetonitrile:water (60:40, v/v) and mobile phase B consisted of isopropanol:acetonitrile:water (88:10:2, v/v/v); both phases contained 10 mM ammonium formate and 0.1% formic acid. Separation was achieved at a flow rate of 260 μL/min using the following solvent gradient: equilibration at 30% B from −3 to 0 min; 30–43% B from 0 to 2 min; 43–55% B from 2 to 2.1 min; 55–65% B from 2.1 to 12 min; 65–85% B from 12 to 18 min; 85–100% B from 18 to 20 min; 100% B from 20 to 25 min; returning to 30% B at 25.1 min and held at 30% B until 28 min.

### Mass spectrometry

LC-MS data were acquired on an Orbitrap Exploris 240 mass spectrometer (Thermo Fisher Scientific) operating with heated electrospray ionization (H-ESI) in both positive and negative ion modes. Source parameters were as follows: spray voltage +3,500 V (positive mode) and -3,000 V (negative mode); sheath gas 40; auxiliary gas 10; sweep gas 1; vaporizer temperature 275°C; ion transfer tube temperature 300°C. Full-scan MS1 data were collected at the orbitrap resolution of 120,000 over a scan range of *m/z* 200–1,700, with a maximum injection time of 50 ms, an RF lens setting of 40%, AGC target set to Standard, and mild trapping enabled. Sub-ppm mass accuracy was maintained throughout data acquisition using the EASY-IC internal calibrant.

### Data-dependent MS2 data acquisition

To increase detection sensitivity, individual samples were used for full MS1 scan and pooled samples were used for quality control (QC) and lipid annotation. QC pools were injected at the beginning of the experiment to equilibrate the column and subsequently every 10-15 samples to correct for instrumental retention time drift over the course of the entire experiment. QC pools were analyzed by data-dependent MS2 acquisition at the orbitrap resolution of 30,000, using a cycle time of 1.5 s. Precursor ions were isolated with a 1.0 *m/z* isolation window and fragmented by higher-energy collisional dissociation (HCD) at stepped normalized collision energies of 25% and 30% for the positive mode and 20%, 30%, and 40% for the negative mode. Dynamic exclusion was applied for 3 s with a mass tolerance of 5 ppm, and the intensity threshold for MS2 triggering was set to 2.0 × 10^4^ counts, the maximum injection time was 54 ms, and the first mass was set to *m/z* 75. Comprehensive fragmentation coverage was achieved using the AcquireX data acquisition workflow (Thermo Fisher Scientific), which employs iterative precursor exclusion across pooled samples (two iterative injections of QC mixes of all conditions plus two injections from individual pools representing individual dietary or RNAi treatment conditions) to prioritize fragmentation of low-abundance and previously unfragmented ions while excluding background and redundant signals.

### Data analysis

Raw LC–MS data were processed and annotated using Compound Discoverer 3.4 with the integrated LipidSearch [MKI] node that used LipidSearch 5.2 (Thermo Fisher Scientific). Data acquired in positive- and negative-ion modes were analyzed independently.

Briefly, Compound Discoverer was used for untargeted peak detection, feature alignment with a retention-time tolerance of ±0.2 min, elemental composition assignment based on accurate mass and isotopic pattern, and compound annotation. Lipid species were annotated in LipidSearch by matching accurate precursor masses (±5 ppm) and experimental MS/MS spectra to database-derived lipid ions and predicted fragment ions. Specifically, candidate annotations were first filtered by precursor mass accuracy and then evaluated based on MS/MS spectral matching to lipid class-specific diagnostic fragments, including headgroup ions, neutral losses, and fatty acyl chain fragments. LipidSearch ranked the top candidate annotations using combined precursor and fragment evidence, enabling assignment of lipid class and fatty acyl composition when supported by the MS/MS data. Top annotations with fatty acyl compositions were filtered using a signal-to-noise ratio threshold of ≥3.0. To increase annotation confidence, we further performed manual validation of individual lipid assignments by inspecting chromatographic peak shape and the correspondence between experimental MS/MS spectra and predicted fragment ions in the LipidSearch database. Lipid annotations are reported using LipidSearch nomenclature; for example, PE(14:0_16:0) denotes a phosphatidylethanolamine species containing 14:0 and 16:0 fatty acyl chains, with acyl chain composition assigned but sn-position unresolved.

Ion abundance data for validated lipid species were exported from Compound Discoverer, and normalization was performed separately for each experiment in Excel using a two-step workflow. First, ion counts were normalized to total protein concentration to account for variation in sample input. For each sample, lipid ion counts were multiplied by a scaling factor equal to the mean protein concentration across all samples within the same experiment divided by the protein concentration of that sample. Second, protein-normalized ion counts were normalized to the corresponding deuterated internal standard from the same lipid class in the equiSPLASH mixture (Avanti Polar Lipids) to account for lipid class-dependent differences in ionization efficiency and ion suppression. For each lipid class, protein-normalized ion counts were multiplied by a scaling factor equal to the mean internal standard ion count across all samples within the same experiment divided by the internal standard ion count in that sample. For example, individual triacylglycerol (TG) species were normalized to the deuterated TG internal standard TG 15:0_18:1_15:0(d7). Internal standards were available for phosphatidylcholine (PC), phosphatidylethanolamine (PE), phosphatidylglycerol (PG), phosphatidylinositol (PI), phosphatidylserine (PS), lysophosphatidylcholine (LPC), lysophosphatidylethanolamine (LPE), monoacylglycerol (MG), diacylglycerol (DG), triacylglycerol (TG), cholesteryl ester (ChE), sphingomyelin (SM), and ceramide (Cer). For lipid classes not represented in the internal standard mixture, including cardiolipin (CL), phosphatidic acid (PA), and N-acyl ethanolamines (NAE), downstream analyses were performed using protein-normalized ion counts. Fold-change differences between experimental and control groups were calculated from fully normalized ion counts when class-matched internal standards were available, or from protein-normalized ion counts when they were not. Normalized ion counts and log2 fold-change data generated in this study are provided in S1 Data and S2 Data.

### Worm lipid supplementation

Bulk lipid extracts from individual *Bifidobacteria* strains were extracted from weight-normalized, concentrated bacterial pellets using the same procedure described in the “Worm lipid extraction for lipidomics” section, without adding the EquiSPLASH lipid standard mix, which was only used for lipidomics. Dried lipid extracts were reconstituted in 75 µL of 100% methanol, followed by the addition of 75 µL of sterile water, then aliquoted and stored at −20°C until the day of supplementation. 50% methanol solvent was used as the vehicle control and was stored under identical conditions. For oxidative stress resistance assays, worms were age-synchronized and maintained as previously described. On the day of use, 150 µL of lipid extract resuspended in 50% methanol or the 50% methanol vehicle control was evenly spread onto the plates and air-dried under a laminar flow hood in the dark. Plates were then seeded with 150 µL of heat-killed *E. coli* and were then air-dried. Synchronized adult day 1 worms were transferred to seeded plates and maintained until adult day 8, at which point they were subjected to paraquat-induced oxidative stress as described above. Worms were transferred every other day, and lipid extracts were freshly applied to the new transfer plates on each day of use. Survival data were plotted as Kaplan–Meier curves using GraphPad Prism, and statistical significance was assessed by log-rank test using the JMP software. Two-way ANOVA was used to evaluate the significance of interactions between experimental conditions. Representative survival curves are presented in the main figures, and summary of survival statistics is provided in S4 Table.

## Supporting information

Compiled document of supporting information, figures, and tables.

## Acknowledgements

We thank the *Caenorhabditis* Genetics Center for providing worm strains; Malene Hansen, Dong Yan, Ryan Baugh, David Tobin, Kenichi Yokoyama, Raphael Valdivia, Dennis Ko, and Violet Beaty for helpful discussions; Brian De’Felice and Xiaoai Zhao for advice on lipidomics liquid chromatography method development. Supported by NIH 5-P30-DK034987-35-39, Mallinckrodt Foundation Grant, Glenn Foundation for Medical Research and AFAR Grant for Junior Faculty, Norins-Benter Award, and Whitehead Scholar (S.H.); Duke University School of Medicine Biomedical PhD Student Research Pilot Grant (Y.L.); and Duke University Dean’s Graduate Fellowship (G.D.T.).

## Author Contributions

Y.L. conceived the study under the guidance of S.H. Y.L. designed and performed all experiments except for the ones described below. G.D.T. performed ROS quantification and pharyngeal pumping assays and assisted with selected oxidative stress assays. S.H. optimized previously published LC-MS/MS methods for the in-house mass spectrometry platform, collected and analyzed lipidomics data, and assisted with selected lifespan and oxidative stress assays. Y.L. and S.H. wrote the manuscript with input from G.D.T.

## Data availability

All study data are included in this manuscript. All lipidomics raw data have been deposited at the NIH Common Fund’s National Metabolomics Data Repository (NMDR) website, the Metabolomics Workbench, https://www.metabolomicsworkbench.org, under study number ST004899 for the *Bifidobacteria* diet dataset and study number ST004900 for the *fat-7* RNAi dataset. These data will be made publicly available upon publication.

