## Supplementary material for "Longevity-promoting human gut *Bifidobacteria* strains require distinct cytoprotective pathways and share dependence on host lipid regulation": Compiled document of supporting information, figures, and tables.

**S1 Fig. Genetic interactions between *Bifidobacteria* and longevity-associated regulators in healthspan and lifespan**. **A**. Lifespan of worms fed *E. coli*, individual pro-longevity *Bifidobacteria* strains, or *E.coli*/*Bifidobacteria* mixtures at 1:1 or 1:4 ratios, with total bacterial biomass held constant across conditions. Bar graph shows percent change in mean lifespan relative to the *E. coli* only control. Each bar represents pooled data from 2–3 replicate plates from one experiment; n ≥ 52 worms per condition. *P* values: two-way ANOVA. N.S., not significant. **B**. None of the three *Bifidobacteria* strains improved survival of middle-aged (adult day 8) worms under acute heat stress at 34°C comparing to the *E. coli* control. Assay was terminated after 12 hours. n ≥ 25 per condition. Not significant for all comparisons by log-rank. **C**. Non-lifespan-extending *Bifidobacteria* strains *B. angulatum* and *B. adolescentis* do not provide protection against paraquat-induced oxidative stress. n ≥ 49. **D-E**. Lifespan extension by *B. breve* is reduced in *nhr-49(nr2041)* (3.95% vs. 26.52% wild-type, **D**) and *daf-16(mu86)* (-0.64% vs. 20.54% wild-type, **E**). *P* < 0.0001 for both pairwise comparisons, two-way ANOVA. *n* ≥ 63 per condition. **F**. RT-qPCR analysis of *sbp-1* transcript levels in worms exposed to empty-vector control or *sbp-1* RNAi bacteria for one day beginning on adult day 1. One-day *sbp-1* RNAi was sufficient to reduce *sbp-1* mRNA levels. Because *sbp-1* RNAi impaired development, RNAi treatment in lifespan assays was initiated on adult day 1, followed by transfer to heat-killed diets on adult day 2. Data are mean ± s.e.m. from 6 biological replicates from one experiment. n > 40 worms per condition. *P* values: two-tailed Mann–Whitney test. **G-I**. The *hlh-30(tm1978)* mutation abolishes lifespan extension by all three *Bifidobacteria* strains and further reduces lifespan below the *E. coli* control levels (-9.83% *B. longum*, -12.31% *B. infantis*, -11.13% *B. breve, P* < 0.0001 by log-rank for all three comparisons). *n* ≥ 22 per condition. **J-M**.

The % lifespan extension by individual *Bifidobacteria* strains relative to *E. coli* control in four long-lived mutants: *eat-2(ad465)*, *raga-1(ok386)*, *rsks-1(ok1255)*, and *rict-1(ft7)*. $n \geq 70$ per condition. *P* values: two-way ANOVA. **N**. The % lifespan extension by *B. longum* or *B. infantis* relative to *E. coli* in two long-lived mutants targeting different subunits of the mitochondrial electron transport chain: *nuo-6(qm200)* (complex I subunit) and *isp-1(qm150)* (complex III subunit). $n \geq 51$ per condition. *P* values: two-way ANOVA. For **N**, day one of adulthood was defined as day one of lifespan due to the extended developmental period of *nuo-6* and *isp-1* mutants relative to wild-type controls. **B, D-E, G-J**, **L-N**, representative of two independent experiments. **A**, **C**, **F**, **K**, from one experiment.

**S2 Fig. Interactions between *Bifidobacteria* diets, host transcription factors, lipid accumulation, and FAT-7. A-B.** Oxidative stress protection conferred by *B. infantis* is not reduced in *daf-16(mu86)* mutants (56.56% vs. 31.42% wild-type, **A**) or *hsf-1(sy441)* (29.42% vs. 25.21% wild-type, **B**). $n \geq 59$ per condition. Representative of two independent experiments. **C**. Oil-Red-O staining intensity in middle-aged (adult day 8) worms fed *B. infantis* or *B. longum* comparing to the *E. coli* controls. Each dot represents one worm; $n > 30$ per condition. Representative of two independent experiments. *P* values: Kruskal-Wallis. **D.** Lifespan extension by *B. infantis* is reduced in *fat-6(tm331);fat-7(wa36)* double mutants (3.50% vs. 17.61% wild-type). $P = 0.005$ by two-way ANOVA. $n \geq 60$ per condition. **E**. RT-qPCR analysis of *fat-5*, *fat-6*, and *fat-7* transcript levels. Mean ± s.e.m from combined data from 2-3 independent experiments, each with 3 biological replicates. *P* values: two-tailed Mann-Whitney with Benjamini-Hochberg correction for multiple comparisons (***P* < 0.01). **F**. RT-qPCR analysis of *fat-7* transcript levels in wild-type and transcription factor-deficient worms fed *B. longum* or *E. coli* control. Mean ±

s.e.m from 2-3 independent experiments, each with 3 biological replicates. *P* values: two-tailed Mann-Whitney with Benjamini-Hochberg correction for multiple comparisons (***P* < 0.01). **G-H.** Lifespan extension by *B. longum* is reduced in *fat-6(tm331);fat-7(wa36)* double mutants (**G**; 4.90% vs. 32.53% wild-type, *P* < 0.0001) and by *fat-7* RNAi (**H**; 4.24% vs. 34.61% empty vector, *P* = 0.0006). *P* values by two-way ANOVA. n ≥ 57 per condition. **I**. Oxidative stress protection by *B. longum* is reduced by *fat-7* RNAi (13.96% vs. 42.38% empty vector). *P* = 0.0135, two-way ANOVA. n ≥ 56 per condition. Representative of two independent experiments.

**S3 Fig. Identification of lipidomic signatures that are *Bifidobacteria*-dependent and longevity-associated. A**. The in-house, LC-MS/MS-based lipidomics tool identifies 450+ lipid species spanning 15 distinct lipid subclasses in the worm lysates. **B**. Principal component analysis separates lipidomic profiles of middle-aged (adult day 8) worms by distinct *Bifidobacteria* diets. The log2 fold-change dataset is normalized to the non-lifespan-extending *B. angulatum* control. **C**-**D**. The lipid species shown represent a panel of *B. infantis*-dependent and longevity-associated that passed the three filtering criteria as described in **Fig 4C**: 1) significantly upregulated by *B. infantis* relative to the non-lifespan-extending *B. angulatum* control (≥ 2-fold, or log2 ≥ 1, adjusted *P* < 0.05), 2) significantly upregulated by *B. infantis* relative to the *E. coli* control using the same threshold, and 3) significantly reduced by *fat-7* RNAi (adjusted *P* < 0.05). Log2 fold-change values are normalized to *B. angulatum* (**C**) or *E. coli* (**D**). **E-H**. Equivalent analysis as in **Fig 4C** identifying *B. longum*-dependent, longevity-associated lipids based on the same three filtering criteria. Log2 fold-change values are normalized to *B. adolescentis* (**E**), *B. angulatum* (**F**), or *E. coli* (**G-H**). Mean ± s.e.m. of one experiment with 5-6 biological replicates per condition, each containing ~500 worms. Lipid species highlighted in red dashed boxes are those shared between the longevity-associated lipid signatures of *B. infantis* (**C**-**D**) and *B. longum* (**E**-**H**). *P* values: two-tailed Mann-Whitney with Benjamini-Hochberg correction for multiple comparisons.

**S1 Table**

Title: List of bacterial strains used in this study

**S2 Table**

Title: List of conserved transcription factors used in this study

**S3 Table**

Title: Summary of lifespan statistics

Additional info: Experiments used in figures are indicated in the right-most column. Mean lifespan and s.e.m. were calculated by log-rank test using triplicate samples, each containing 25-35 worms. # worms: total number of worms died in this assay/(total died + total censored). Censored worms included ones underwent "bagging", exhibited ruptured vulva, or crawled off the plates.

**S4 Table**

Title: Summary of oxidative stress survival statistics

Additional info: Experiments used in figures are indicated in the right-most column. Survival was scored hourly for up to 12 hours. Worms that were still alive at the end of experiment were censored. Mean lifespan and s.e.m. were calculated by log-rank test using duplicate samples, each containing 25-35 worms. # worms: total number of worms died in this assay/(total died + total censored). Censored worms were defined as those still alive at the 12-hr timepoint of experiment termination.

**S5 Table**

Title: List of worm strains used in this study

**S6 Table**

Title: List of RT-qPCR primers used in this study

**S1 Data**

Title: *Bifidobacteria* diet lipidomics data

Worksheet: study_metadata

| | |
|---|---|
| Sample_ID | sample IDs used in the remaining worksheets |
| Detection_method | reverse phase c18 positive or c18 negative methods |
| Sample_source | *C. elegans* for all samples |
| Genotype | wild-type for all samples |
| Treatment | Worms were fed live *E. coli* from hatching through day 1 of reproductive adulthood, then switched to heat-killed *E. coli* or individual *Bifidobacteria* diets until sample collection on day 8 of adulthood. |

Worksheet: lipid_metadata

This worksheet contains details on individual annotated lipids in this dataset, including chemical name, molecular formula, *m/z*, reference ion, retention time in minutes, and detection method.

Worksheets: c18neg_norm_ion_count and c18pos_norm_ion_count

Normalized ion counts are reported for individual lipids detected by reverse-phase C18 LC-MS/MS in negative or positive ion mode. Raw ion counts were normalized to protein concentration and class-specific lipid internal standards as described in Methods. Sample IDs correspond to the metadata provided in "study_metadata."

Worksheets: log2_fc_B_adolescentis and log2_fc_B_angulatum

Log2 fold change values were calculated from normalized ion counts by comparing lifespan-extending *Bifidobacteria* strains (*B. longum*, *B. infantis*, and *B. breve*) with non-lifespan-extending control strains. Values in "log2_fc_B_adolescentis" were calculated relative to *B. adolescentis*, and values in "log2_fc_B_angulatum" were calculated relative to *B. angulatum*.

**S2 Data**

Title: *fat-7* RNAi lipidomics data

Worksheet: study_metadata

| | |
|---|---|
| Sample_ID | sample IDs used in the remaining worksheets |
| Detection_method | reverse phase c18 positive or c18 negative methods |
| Sample_source | *C. elegans* for all samples |
| Genotype | wild-type for all samples |

| | |
|---|---|
| Treatment | Worms were fed live *E. coli* carrying either empty vector or *fat-7* RNAi from hatching through day 1 of reproductive adulthood, then shifted to heat-killed *E. coli*, *B. infantis*, or *B. longum* diets until sample collection on day 8 of adulthood. |

Worksheet: lipid_metadata

This worksheet contains details on individual annotated lipids in this dataset, including chemical name, molecular formula, *m/z*, reference ion, retention time in minutes, and detection method.

Worksheets: c18neg_norm_ion_count and c18pos_norm_ion_count

Normalized ion counts are reported for individual lipids detected by reverse-phase C18 LC-MS/MS in negative or positive ion mode. Raw ion counts were normalized to protein concentration and class-specific lipid internal standards as described in Methods. Sample IDs correspond to the metadata provided in "study_metadata."

Worksheet: log2_fc_Empty_vector

Log2 fold change values were calculated from normalized ion counts by comparing individual conditions (empty vector + *B. longum*, empty vector + *B. infantis*, *fat-7* RNAi + *B. longum*, *fat-7* RNAi + *B. infantis*, and *fat-7* RNAi + *E. coli*) with the control diet of empty vector + *E. coli.*

### S1 Figure

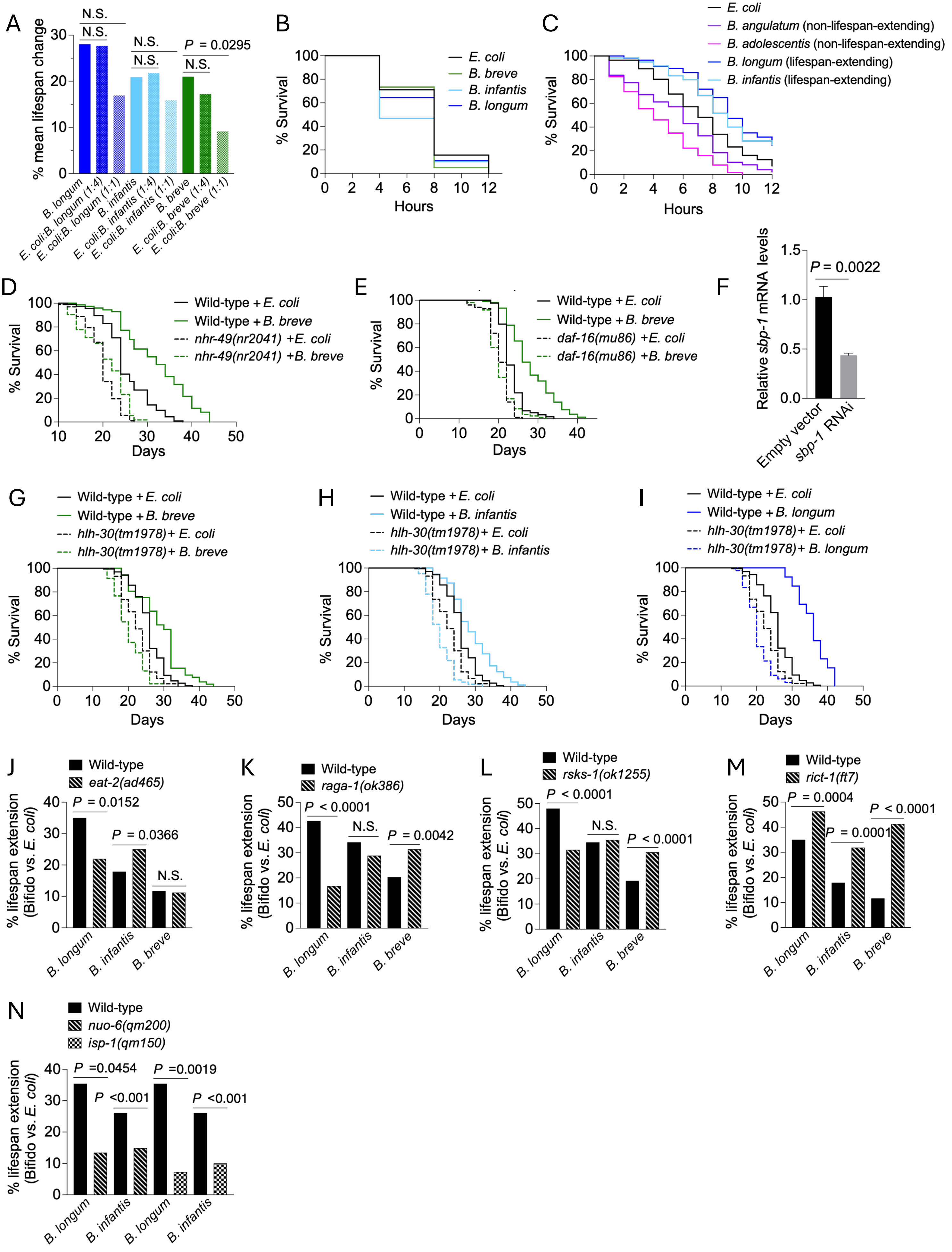

A
% mean lifespan change
N.S.
N.S.
N.S.
N.S.
P = 0.0295
N.S.
B. longum
E. coli:B. longum (1:4)
E. coli:B. longum (1:1)
B. infantis
E. coli:B. infantis (1:4)
E. coli:B. infantis (1:1)
B. breve
E. coli:B. breve (1:4)
E. coli:B. breve (1:1)
B
% Survival
Hours
E. coli
B. breve
B. infantis
B. longum
C
% Survival
Hours
E. coli
B. angulatum (non-lifespan-extending)
B. adolescentis (non-lifespan-extending)
B. longum (lifespan-extending)
B. infantis (lifespan-extending)
D
% Survival
Days
Wild-type + E. coli
Wild-type + B. breve
nhr-49(nr2041) +E. coli
nhr-49(nr2041) +B. breve
E
% Survival
Days
Wild-type + E. coli
Wild-type + B. breve
daf-16(mu86) +E. coli
daf-16(mu86) +B. breve
F
Relative sbp-1 mRNA levels
P = 0.0022
Empty vector
sbp-1 RNAi
G
Wild-type + E. coli
Wild-type + B. breve
hlh-30(tm1978)+ E. coli
hlh-30(tm1978)+ B. breve
% Survival
Days
H
Wild-type + E. coli
Wild-type + B. infantis
hlh-30(tm1978)+ E. coli
hlh-30(tm1978)+ B. infantis
% Survival
Days
I
Wild-type + E. coli
Wild-type + B. longum
hlh-30(tm1978)+ E. coli
hlh-30(tm1978)+ B. longum
% Survival
Days
J
Wild-type
eat-2(ad465)
% lifespan extension (Bifido vs. E. coli)
P = 0.0152
P = 0.0366
N.S.
B. longum
B. infantis
B. breve
K
Wild-type
raga-1(ok386)
% lifespan extension (Bifido vs. E. coli)
P < 0.0001
N.S.
P = 0.0042
B. longum
B. infantis
B. breve
L
Wild-type
rsks-1(ok1255)
% lifespan extension (Bifido vs. E. coli)
P < 0.0001
N.S.
P < 0.0001
B. longum
B. infantis
B. breve
M
Wild-type
rict-1(ft7)
% lifespan extension (Bifido vs. E. coli)
P = 0.0004
P = 0.0001
P < 0.0001
B. longum
B. infantis
B. breve
N
Wild-type
nuo-6(qm200)
isp-1(qm150)
% lifespan extension (Bifido vs. E. coli)
P =0.0454
P <0.001
P =0.0019
P <0.001
B. longum
B. infantis
B. longum
B. infantis

### S2 Figure

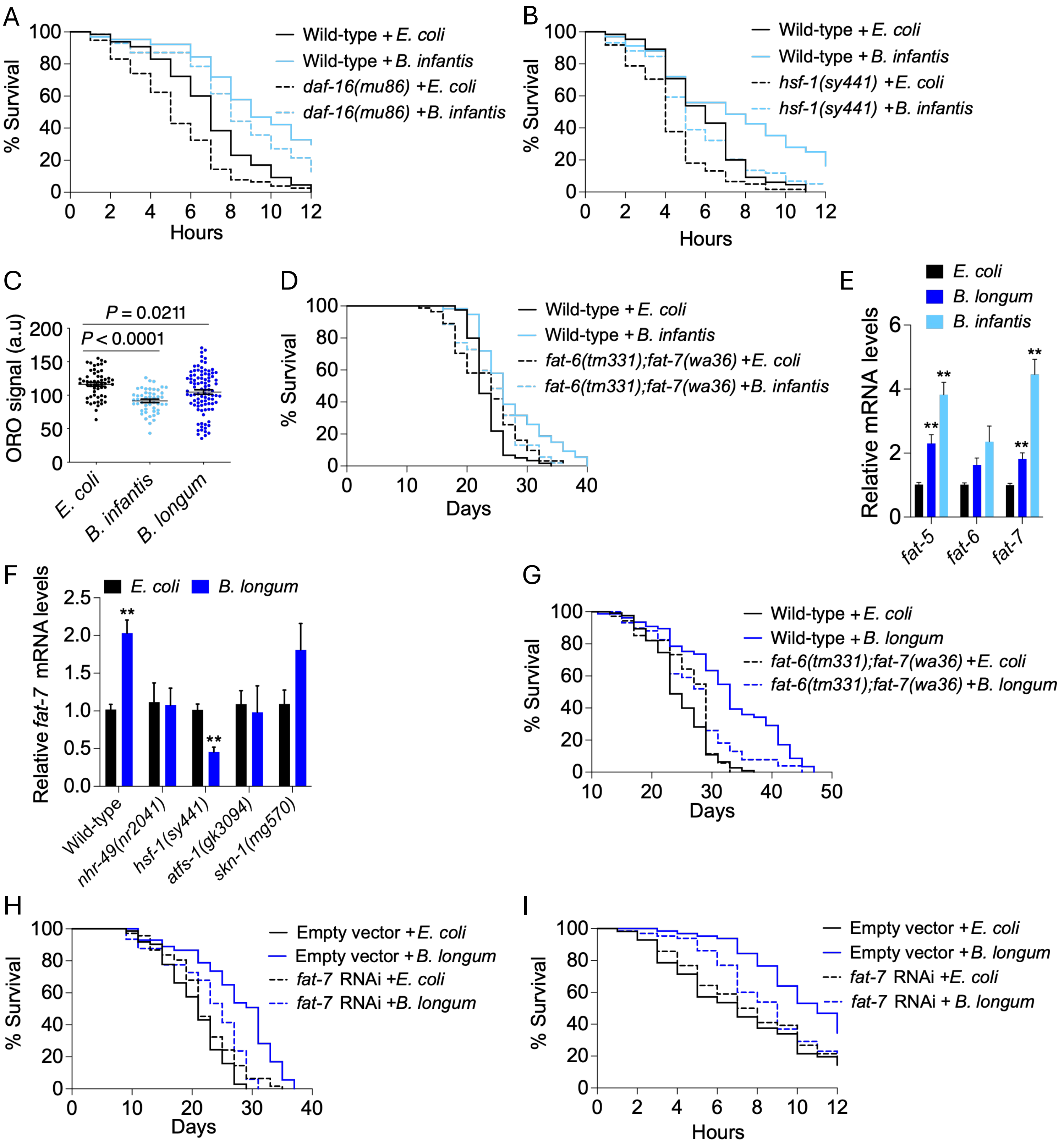

A
% Survival
Hours
Wild-type + E. coli
Wild-type + B. infantis
daf-16(mu86) + E. coli
daf-16(mu86) + B. infantis
B
% Survival
Hours
Wild-type + E. coli
Wild-type + B. infantis
hsf-1(sy441) + E. coli
hsf-1(sy441) + B. infantis
C
ORO signal (a.u)
P = 0.0211
P < 0.0001
E. coli
B. infantis
B. longum
D
% Survival
Days
Wild-type + E. coli
Wild-type + B. infantis
fat-6(tm331);fat-7(wa36) + E. coli
fat-6(tm331);fat-7(wa36) + B. infantis
E
Relative mRNA levels
E. coli
B. longum
B. infantis
fat-5
fat-6
fat-7
F
Relative fat-7 mRNA levels
E. coli
B. longum
Wild-type
nhr-49(nr2041)
hsf-1(sy441)
atfs-1(gk3094)
skn-1(mg570)
G
% Survival
Days
Wild-type + E. coli
Wild-type + B. longum
fat-6(tm331);fat-7(wa36) + E. coli
fat-6(tm331);fat-7(wa36) + B. longum
H
% Survival
Days
Empty vector + E. coli
Empty vector + B. longum
fat-7 RNAi + E. coli
fat-7 RNAi + B. longum
I
% Survival
Hours
Empty vector + E. coli
Empty vector + B. longum
fat-7 RNAi + E. coli
fat-7 RNAi + B. longum

S3 Figure

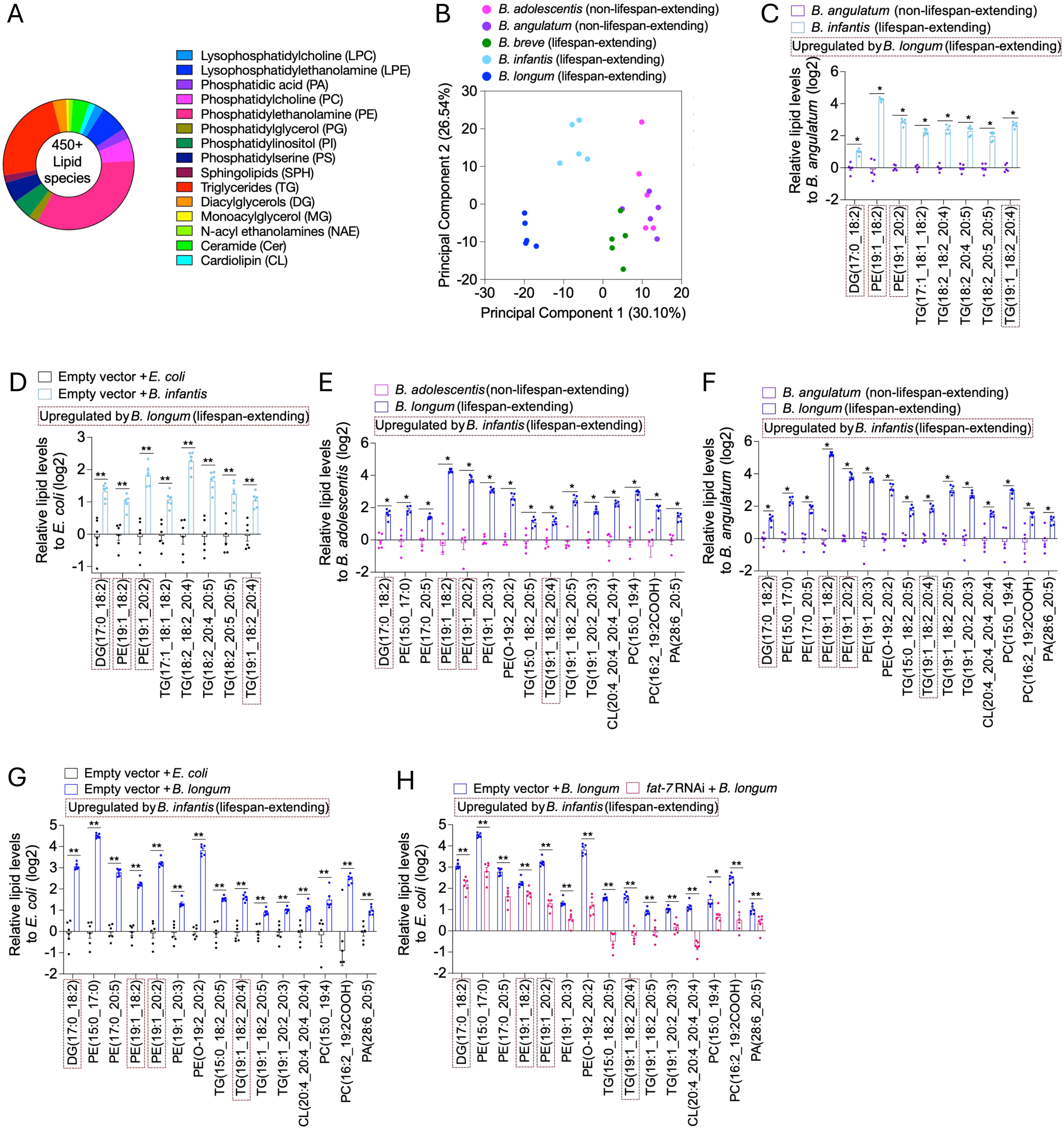
A
450+ Lipid species
Lysophosphatidylcholine (LPC)
Lysophosphatidylethanolamine (LPE)
Phosphatidic acid (PA)
Phosphatidylcholine (PC)
Phosphatidylethanolamine (PE)
Phosphatidylglycerol (PG)
Phosphatidylinositol (PI)
Phosphatidylserine (PS)
Sphingolipids (SPH)
Triglycerides (TG)
Diacylglycerols (DG)
Monoacylglycerol (MG)
N-acyl ethanolamines (NAE)
Ceramide (Cer)
Cardiolipin (CL)
B
B. adolescentis (non-lifespan-extending)
B. angulatum (non-lifespan-extending)
B. breve (lifespan-extending)
B. infantis (lifespan-extending)
B. longum (lifespan-extending)
Principal Component 1 (30.10%)
Principal Component 2 (26.54%)
C
B. angulatum (non-lifespan-extending)
B. infantis (lifespan-extending)
Upregulated by B. longum (lifespan-extending)
Relative lipid levels to B. angulatum (log2)
D
Empty vector + E. coli
Empty vector + B. infantis
Upregulated by B. longum (lifespan-extending)
Relative lipid levels to E. coli (log2)
E
B. adolescentis (non-lifespan-extending)
B. longum (lifespan-extending)
Upregulated by B. infantis (lifespan-extending)
Relative lipid levels to B. adolescentis (log2)
F
B. angulatum (non-lifespan-extending)
B. longum (lifespan-extending)
Upregulated by B. infantis (lifespan-extending)
Relative lipid levels to B. angulatum (log2)
G
Empty vector + E. coli
Empty vector + B. longum
Upregulated by B. infantis (lifespan-extending)
Relative lipid levels to E. coli (log2)
H
Empty vector + B. longum
fat-7 RNAi + B. longum
Upregulated by B. infantis (lifespan-extending)
Relative lipid levels to E. coli (log2)
DG(17:0_18:2)
PE(15:0_17:0)
PE(17:0_20:5)
PE(19:1_18:2)
PE(19:1_20:2)
PE(19:1_20:3)
PE(O-19:2_20:2)
TG(15:0_18:2_20:5)
TG(17:1_18:1_18:2)
TG(18:2_18:2_20:4)
TG(18:2_20:4_20:5)
TG(18:2_20:5_20:5)
TG(19:1_18:2_20:4)
TG(19:1_18:2_20:5)
TG(19:1_20:2_20:3)
CL(20:4_20:4_20:4)
PC(15:0_19:4)
PC(16:2_19:2COOH)
PA(28:6_20:5)

**S1 Table: List of bacterial strains used in this study**

| **Strain name** |
|---|
| *Escherichia coli* OP50 |
| *Escherichia coli* HT115 |
| *Bifidobacterium adolescentis* L2-32 |
| *Bifidobacterium angulatum* DSMZ 20098 |
| *Bifidobacterium bifidum* DSMZ 20456 |
| *Bifidobacterium breve* DSMZ 20213 |
| *Bifidobacterium breve* UCC2003 |
| *Bifidobacterium catenulatum* DSMZ 16992 |
| *Bifidobacterium dentium* ATCC 27678 |
| *Bifidobacterium infantis* ATCC 15697 |
| *Bifidobacterium longum* NCC 2705 |
| *Bifidobacterium longum* reuter ATCC 55813 |
| *Bifidobacterium pseudocatenulatum* DSMZ 20438 |

**S2 Table: List of conserved transcription factors**

| *C. elegans* | Mammalian functional ortholog |
|---|---|
| ATFS-1 | ATF5 |
| HSF-1 | HSF1 |
| SKN-1 | NRF2 |
| DAF-16 | FOXO |
| NHR-49 | PPARα |
| NHR-80 | HNF4α |
| XBP-1 | XBP1 |
| SBP-1 | SREBP1 |
| HLH-30 | TFEB |

**S3 Table: Summary of lifespan statistics**

| Worm strains | Treatments | Mean +/- s.e.m. (since hatching unless otherwise noted) | *P* values (log-rank) | % lifespan change (compared to the control diet, unless otherwise noted) | # worms (total number of worms died in this assay/(total died + total censored) | Figures |
|---|---|---|---|---|---|---|
| wild-type (N2) | *E. coli* OP50 (control) | 20.9443 +/- 0.4471 | | | 96/118 | Fig 1B |
| | *B. adolescentis* L2-32 | 20.5351 +/- 0.71945 | 0.7581 | -1.95 | 94/110 | Fig 1B |
| | *B. bifidum* DSMZ 20456 | 26.5785 +/- 0.81211 | <0.0001 | 26.90 | 69/92 | Fig 1B |
| | *B. breve* UCC2003 | 25.3813 +/- 0.63306 | <0.0001 | 21.18 | 86/108 | Fig 1B |
| | *B. dentium* ATCC 27678 | 23.1258 +/- 0.71879 | 0.0062 | 10.42 | 79/94 | Fig 1B |
| | *B. longum reuter* ATCC 55813 | 25.606 +/- 0.75621 | <0.0001 | 22.26 | 63/83 | Fig 1B |
| | *B. longum* NCC 2705 | 28.9037 +/- 0.68096 | <0.0001 | 38.00 | 71/99 | Fig 1B |
| | *B. pseudocatenulatum* DSMZ 20438 | 25.8167 +/- 0.713 | <0.0001 | 23.26 | 80/104 | Fig 1B |
| wild-type (N2) | *E. coli* OP50 (control) | 21.4779 +/- 0.44952 | | | 89/107 | Fig 1B |
| | *B. catenulatum* DSMZ 16992 | 18.9244 +/- 0.51179 | 0.0002 | -11.89 | 84/104 | Fig 1B |
| | *B. longum* NCC 2705 | 29.6669 +/- 0.71914 | <0.0001 | 38.13 | 89/114 | Fig 1B |
| wild-type (N2) | *E. coli* OP50 (control) | 21.3952 +/- 0.40636 | | | 106/118 | Fig 1B |
| | *B. adolescentis* L2-32 | 22.4614 +/- 0.51256 | 0.0517 | 4.98 | 113/127 | Fig 1B |
| | *B. angulatum* DSMZ 20098 | 20.4231 +/- 0.44064 | 0.0497 | -4.54 | 116/130 | Fig 1B |
| | *B. breve* UCC2003 | 24.2618 +/- 0.50068 | <0.0001 | 13.40 | 98/124 | Fig 1B |
| | *B. catenulatum* DSMZ 16992 | 18.0355 +/- 0.50954 | <0.0001 | -15.70 | 94/98 | Fig 1B |
| | *B. longum reuter* ATCC 55813 | 24.2244 +/- 0.57754 | <0.0001 | 13.22 | 99/124 | Fig 1B |
| wild-type (N2) | *E. coli* OP50 (control) | 24.5742 +/- 0.43711 | | | 67/83 | Fig 1B |
| | *B. dentium* ATCC 27678 | 27.6076 +/- 0.61331 | <0.0001 | 16.41 | 66/88 | Fig 1B |
| wild-type (N2) | *E. coli* OP50 (control) | 24.2813 +/- 0.52445 | | | 96/107 | Fig 1B |
| | *B. adolescentis* L2-32 | 22.6207 +/- 0.66222 | 0.0751 | -6.84 | 83/103 | Fig 1B |
| | *B. angulatum* DSMZ 20098 | 23.2341 +/- 0.71767 | 0.1826 | -4.31 | 81/103 | Fig 1B |
| | *B. bifidum* DSMZ 20456 | 28 +/- 0.63232 | <0.0001 | 15.32 | 73/100 | Fig 1B |
| | *B. pseudocatenulatum* DSMZ 20438 | 30.1481 +/- 1.68511 | <0.0001 | 24.16 | 33/60 | Fig 1B |
| wild-type (N2) | *E. coli* OP50 (control) | 24.6723 +/- 0.33574 | | | 89/111 | Fig 1B, 1C, 2A, 2F, S1L |
| | *B. breve* DSMZ 20213 | 29.4299 +/- 0.83979 | <0.0001 | 19.28 | 60/87 | Fig 1B, 1C, 2A, S1L |
| | *B. infantis* ATCC 15697 | 33.2031 +/- 0.89139 | <0.0001 | 34.58 | 75/100 | Fig 1B, 1C, 2A, 2F, S1L |
| | *B. longum* NCC 2705 | 36.5162 +/- 0.86566 | <0.0001 | 48.00 | 57/99 | Fig 1C, 2A, 2F, S1L |
| *skn-1(mg570)* | *E. coli* OP50 | 25.1705 +/- 0.48034 | | | 110/128 | Fig 2A, 2F |
| | *B. breve* DSMZ 20213 | 29.9483 +/- 0.74738 | <0.0001 (compared to *skn-1(mg570)* + *E. coli* OP50) | 18.98 (compared to *skn-1(mg570)* + *E. coli* OP50) | 65/105 | Fig 2A |
| | *B. infantis* ATCC 15697 | 30.7044 +/- 0.6165 | <0.0001 (compared to *skn-1(mg570)* + *E. coli* OP50) | 21.99 (compared to *skn-1(mg570)* + *E. coli* OP50) | 87/121 | Fig 2A, 2F |
| | *B. longum* NCC 2705 | 31.4671 +/- 0.86847 | <0.0001 (compared to *skn-1(mg570)* + *E. coli* OP50) | 25.02 (compared to *skn-1(mg570)* + *E. coli* OP50) | 64/104 | Fig 2A, 2F |
| *eat-2(ad465)* | *E. coli* OP50 | 28.0923 +/- 0.49668 | | | 100/124 | |
| | *B. breve* DSMZ 20213 | 33.8147 +/- 0.56671 | <0.0001 (compared to *eat-2(ad465)* + *E. coli* OP50) | 20.37 (compared to *eat-2(ad465)* + *E. coli* OP50) | 81/110 | |
| | *B. longum* NCC 2705 | 37.2023 +/- 0.77682 | <0.0001 (compared to *eat-2(ad465)* + *E. coli* OP50) | 32.43 (compared to *eat-2(ad465)* + *E. coli* OP50) | 83/108 | |
| *rsks-1(ok1255)* | *E. coli* OP50 | 33.5265 +/- 0.47531 | | | 86/99 | Fig S1L |
| | *B. breve* DSMZ 20213 | 43.8041 +/- 0.61603 | <0.0001 (compared to *rsks-1(ok1255)* + *E. coli* OP50) | 30.66 (compared to *rsks-1(ok1255)* + *E. coli* OP50) | 76/107 | Fig S1L |
| | *B. infantis* ATCC 15697 | 45.445 +/- 0.57014 | <0.0001 (compared to *rsks-1(ok1255)* + *E. coli* OP50) | 35.55 (compared to *rsks-1(ok1255)* + *E. coli* OP50) | 84/93 | Fig S1L |
| | *B. longum* NCC 2705 | 44.1154 +/- 0.564 | <0.0001 (compared to *rsks-1(ok1255)* + *E. coli* OP50) | 31.58 (compared to *rsks-1(ok1255)* + *E. coli* OP50) | 62/85 | Fig S1L |

| Worm strains | Treatments | Mean +/- s.e.m. (since hatching unless otherwise noted) | *P* values (log-rank) | % lifespan change (compared to the control diet, unless otherwise noted) | # worms (total number of worms died in this assay/(total died + total censored) | Figures |
|---|---|---|---|---|---|---|
| wild-type (N2) | *E. coli* OP50 (control) | 20.5838 +/- 0.4606 | | | 97/103 | Fig S1A |
| | *B. longum* NCC 2705 | 26.3575 +/- 0.9893 | <0.0001 | 28.05 | 46/52 | Fig S1A |
| | 1:4 (*E. coli* OP50:*B. longum* NCC 2705) | 26.2795 +/- 1.0019 | <0.0001 | 27.67 | 55/65 | Fig S1A |
| | 1:1 (*E. coli* OP50:*B. longum* NCC 2705) | 24.0657 +/- 0.5476 | <0.0001 | 16.92 | 83/89 | Fig S1A |
| | *B. infantis* ATCC 15697 | 24.8944 +/- 0.6602 | <0.0001 | 20.94 | 61/64 | Fig S1A |
| | 1:4 (*E. coli* OP50:*B. infantis* ATCC 15697) | 25.0769 +/- 0.8359 | <0.0001 | 21.83 | 77/84 | Fig S1A |
| | 1:1 (*E. coli* OP50:*B. infantis* ATCC 15697) | 23.8568 +/- 0.7025 | <0.0001 | 15.90 | 89/93 | Fig S1A |
| | *B. breve* DSMZ 20213 | 24.908 +/- 0.7421 | <0.0001 | 21.01 | 52/61 | Fig S1A |
| | 1:4 (*E. coli* OP50:*B. breve* DSMZ 20213) | 24.128 +/- 0.6558 | <0.0001 | 17.22 | 99/104 | Fig S1A |
| | 1:1 (*E. coli* OP50:*B. breve* DSMZ 20213) | 22.4728 +/- 0.5667 | 0.0054 | 9.18 | 96/102 | Fig S1A |
| wild-type (N2) | *E. coli* OP50 (control) | 24.1218 +/- 0.38151 | | | 93/97 | Fig 2A, 2B, 3G |
| | *B. breve* DSMZ 20213 | 27.9275 +/- 1.96415 | 0.0028 | 15.78 | 19/27 | |
| | *B. infantis* ATCC 15697 | 30.1931 +/- 0.73414 | <0.0001 | 25.17 | 82/106 | Fig 2A, 2B, 3G |
| | *B. longum* NCC 2705 | 33.2137 +/- 0.73116 | <0.0001 | 37.69 | 68/89 | Fig 2A, 2B |
| *nhr-49(nr2041)* | *E. coli* OP50 | 20.2843 +/- 0.38147 | | | 86/92 | Fig 2A, 2B |
| | *B. infantis* ATCC 15697 | 19.1767 +/- 0.34547 | 0.0465 (compared to *nhr-49(nr2041)* + *E. coli* OP50) | -5.46 (compared to *nhr-49(nr2041)* + *E. coli* OP50) | 101/109 | Fig 2A, 2B |
| | *B. longum* NCC 2705 | 23.6376 +/- 0.68358 | <0.0001 (compared to *nhr-49(nr2041)* + *E. coli* OP50) | 16.53 (compared to *nhr-49(nr2041)* + *E. coli* OP50) | 71/79 | Fig 2A, 2B |
| *fat-5(tm420)* | *E. coli* OP50 | 23.7215 +/- 0.34163 | | | 110/126 | Fig 3G |
| | *B. infantis* ATCC 15697 | 28.4793 +/- 0.70209 | <0.0001 (compared to *fat-5(tm420)* + *E. coli* OP50) | 20.06 (compared to *fat-5(tm420)* + *E. coli* OP50) | 85/92 | Fig 3G |
| | *B. longum* NCC 2705 | 35.4717 +/- 0.9446 | <0.0001 (compared to *fat-5(tm420)* + *E. coli* OP50) | 49.53 (compared to *fat-5(tm420)* + *E. coli* OP50) | 59/79 | |
| wild-type (N2) | *E. coli* OP50 (control) | 25.2682 +/- 0.48564 | | | 113/117 | Fig 2A, 2C, S1D |
| | *B. breve* DSMZ 20213 | 31.9701 +/- 0.93624 | <0.0001 | 26.52 | 62/76 | Fig 2A, S1D |
| | *B. infantis* ATCC 15697 | 35.5012 +/- 0.98218 | <0.0001 | 40.50 | 60/81 | Fig 2A |
| | *B. longum* NCC 2705 | 35.5955 +/- 0.79662 | <0.0001 | 40.87 | 53/72 | Fig 2A, 2C |
| *daf-16(mu86)* | *E. coli* OP50 | 20.8933 +/- 0.29711 | | | 107/118 | Fig 2A, 2C |
| | *B. longum* NCC 2705 | 28.9308 +/- 0.66079 | <0.0001 (compared to *daf-16(mu86)* + *E. coli* OP50) | 38.47 (compared to *daf-16(mu86)* + *E. coli* OP50) | 63/89 | Fig 2A, 2C |
| *hsf-1(sy441)* | *E. coli* OP50 | 22.3682 +/- 0.31188 | | | 94/107 | Fig 2A |
| | *B. infantis* ATCC 15697 | 23.8606 +/- 0.58199 | 0.0006 (compared to *hsf-1(sy441)* + *E. coli* OP50) | 6.67 (compared to *hsf-1(sy441)* + *E. coli* OP50) | 76/79 | Fig 2A |
| *nhr-49(nr2041)* | *E. coli* OP50 | 19.819 +/- 0.3554 | | | 96/98 | Fig 2A, S1D |
| | *B. breve* DSMZ 20213 | 20.602 +/- 0.64138 | 0.0095 (compared to *nhr-49(nr2041)* + *E. coli* OP50) | 3.95 (compared to *nhr-49(nr2041)* + *E. coli* OP50) | 60/63 | Fig 2A, S1D |
| | *B. longum* NCC 2705 | 23.9286 +/- 0.60156 | <0.0001 (compared to *nhr-49(nr2041)* + *E. coli* OP50) | 20.74 (compared to *nhr-49(nr2041)* + *E. coli* OP50) | 62/72 | Fig 2A |
| wild-type (N2) | *E. coli* OP50 (control) | 25.5122 +/- 0.5155 | | | 84/90 | Fig 2A, 2C, 3G |
| | *B. breve* DSMZ 20213 | 28.5998 +/- 1.01559 | 0.0024 | 12.10 | 33/41 | Fig 2A |
| | *B. infantis* ATCC 15697 | 29.6227 +/- 1.5232 | <0.0001 | 16.11 | 24/26 | Fig 2A, 2C, 3G |
| | *B. longum* NCC 2705 | 31.716 +/- 0.84191 | <0.0001 | 24.32 | 56/65 | Fig 2A |
| *nhr-80(tm1011)* | *E. coli* OP50 | 21.1091 +/- 0.65243 | | | 61/63 | Fig 2A |
| | *B. breve* DSMZ 20213 | 26.9348 +/- 1.02109 | <0.0001 (compared to *nhr-80(tm1011)* + *E. coli* OP50) | 27.60 (compared to *nhr-80(tm1011)* + *E. coli* OP50) | 58/60 | Fig 2A |
| | *B. infantis* ATCC 15697 | 25.5556 +/- 0.91178 | <0.0001 (compared to *nhr-80(tm1011)* + *E. coli* OP50) | 21.06 (compared to *nhr-80(tm1011)* + *E. coli* OP50) | 72/72 | Fig 2A |
| | *B. longum* NCC 2705 | 27.353 +/- 0.96775 | <0.0001 (compared to *nhr-80(tm1011)* + *E. coli* OP50) | 29.58 (compared to *nhr-80(tm1011)* + *E. coli* OP50) | 58/59 | Fig 2A |

| Worm strains | Treatments | Mean +/- s.e.m. (since hatching unless otherwise noted) | *P* values (log-rank) | % lifespan change (compared to the control diet, unless otherwise noted) | # worms (total number of worms died in this assay/(total died + total censored) | Figures |
|---|---|---|---|---|---|---|
| *daf-16(mu86)* | *E. coli* OP50 | 18.9261 +/- 0.38328 | | | 106/110 | Fig 2A, 2C |
| | *B. breve* DSMZ 20213 | 19.3333 +/- 0.61095 | 0.4048 (compared to *daf-16(mu86)* + *E. coli* OP50) | 2.15 (compared to *daf-16(mu86)* + *E. coli* OP50) | 60/60 | Fig 2A |
| | *B. infantis* ATCC 15697 | 19.6775 +/- 0.54436 | 0.2211 (compared to *daf-16(mu86)* + *E. coli* OP50) | 3.97 (compared to *daf-16(mu86)* + *E. coli* OP50) | 69/74 | Fig 2A, 2C |
| *fat-7(wa36)* | *E. coli* OP50 | 24.8076 +/- 0.6697 | | | 74/79 | Fig 3G |
| | *B. infantis* ATCC 15697 | 27.849 +/- 0.76961 | 0.0106 (compared to *fat-7(wa36)* + *E. coli* OP50) | 12.26 (compared to *fat-7(wa36)* + *E. coli* OP50) | 47/60 | Fig 3G |
| | *B. longum* NCC 2705 | 31.8288 +/- 0.79674 | <0.0001 (compared to *fat-7(wa36)* + *E. coli* OP50) | 28.30 (compared to *fat-7(wa36)* + *E. coli* OP50) | 62/82 | |
| wild-type (N2) | *E. coli* OP50 (control) | 24.8831 +/- 0.41805 | | | 94/99 | Fig 2A, 2D, 2E |
| | *B. breve* DSMZ 20213 | 29.1322 +/- 1.00715 | <0.0001 | 17.08 | 41/51 | Fig 2A |
| | *B. infantis* ATCC 15697 | 32.5171+/- 0.83743 | <0.0001 | 30.68 | 83/106 | Fig 2A, 2D, 2E |
| | *B. longum* NCC 2705 | 34.9308 +/- 1.03315 | <0.0001 | 40.38 | 53/73 | Fig 2A, 2D |
| *skn-1(mg570)* | *E. coli* OP50 | 27.0511 +/- 0.59345 | | | 86/95 | Fig 2A |
| | *B. breve* DSMZ 20213 | 31.8193 +/- 0.8008 | <0.0001 (compared to *skn-1(mg570)* + *E. coli* OP50) | 17.63 (compared to *skn-1(mg570)* + *E. coli* OP50) | 62/84 | Fig 2A |
| | *B. infantis* ATCC 15697 | 32.6189 +/- 0.77052 | <0.0001 (compared to *skn-1(mg570)* + *E. coli* OP50) | 20.58 (compared to *skn-1(mg570)* + *E. coli* OP50) | 73/85 | Fig 2A |
| | *B. longum* NCC 2705 | 33.4493 +/- 0.82735 | <0.0001 (compared to *skn-1(mg570)* + *E. coli* OP50) | 23.65 (compared to *skn-1(mg570)* + *E. coli* OP50) | 71/82 | Fig 2A |
| *atfs-1(gk3094)* | *E. coli* OP50 | 23.5509 +/- 0.38277 | | | 88/95 | Fig 2A, 2E |
| | *B. infantis* ATCC 15697 | 26.6197 +/- 0.64358 | <0.0001 (compared to *atfs-1(gk3094)* + *E. coli* OP50) | 13.03 (compared to *atfs-1(gk3094)* + *E. coli* OP50) | 63/78 | Fig 2A, 2E |
| *hsf-1(sy441)* | *E. coli* OP50 | 21.206 +/- 0.39106 | | | 81/85 | Fig 2A, 2D |
| | *B. breve* DSMZ 20213 | 24.7845 +/- 0.60103 | <0.0001 (compared to *hsf-1(sy441)* + *E. coli* OP50) | 16.95 (compared to *hsf-1(sy441)* + *E. coli* OP50) | 70/73 | Fig 2A |
| | *B. infantis* ATCC 15697 | 24.9129 +/- 0.55918 | <0.0001 (compared to *hsf-1(sy441)* + *E. coli* OP50) | 17.56 (compared to *hsf-1(sy441)* + *E. coli* OP50) | 77/79 | Fig 2A, 2D |
| | *B. longum* NCC 2705 | 25.4643 +/- 0.65711 | <0.0001 (compared to *hsf-1(sy441)* + *E. coli* OP50) | 20.18 (compared to *hsf-1(sy441)* + *E. coli* OP50) | 56/58 | Fig 2A, 2D |
| *nhr-49(nr2041)* | *E. coli* OP50 | 17.7016 +/- 0.3501 | | | 67/71 | Fig 2A |
| | *B. infantis* ATCC 15697 | 18.6698 +/- 0.78022 | 0.0938 (compared to *nhr-49(nr2041)* + *E. coli* OP50) | 5.47 (compared to *nhr-49(nr2041)* + *E. coli* OP50) | 31/32 | Fig 2A |
| wild-type (N2) | *E. coli* OP50 (control) | 24.0684 +/- 0.68004 | | | 68/87 | Fig 1B, 2A, 2E, S1K |
| | *B. breve* DSMZ 20213 | 28.9392 +/- 0.90513 | <0.0001 | 20.24 | 55/76 | Fig 1B, 2A, S1K |
| | *B. infantis* ATCC 15697 | 32.2825 +/- 0.8025 | <0.0001 | 34.13 | 80/122 | Fig 1B, S1K |
| | *B. longum* NCC 2705 | 34.3332 +/- 0.7179 | <0.0001 | 42.65 | 51/83 | Fig 2A, 2E, S1K |
| *atfs-1(gk3094)* | *E. coli* OP50 | 23.4832 +/- 0.74676 | | | 41/73 | Fig 2A, 2E |
| | *B. breve* DSMZ 20213 | 30.8144 +/- 0.83161 | <0.0001 (compared to *atfs-1(gk3094)* + *E. coli* OP50) | 31.22 (compared to *atfs-1(gk3094)* + *E. coli* OP50) | 51/71 | Fig 2A |
| | *B. longum* NCC 2705 | 28.9107 +/- 0.70852 | <0.0001 (compared to *atfs-1(gk3094)* + *E. coli* OP50) | 23.11 (compared to *atfs-1(gk3094)* + *E. coli* OP50) | 50/72 | Fig 2A, 2E |
| *raga-1(ok386)* | *E. coli* OP50 | 36.8345 +/- 0.83674 | | | 49/70 | Fig S1K |
| | *B. breve* DSMZ 20213 | 48.41 +/- 0.84478 | <0.0001 (compared to *raga-1(ok386)* + *E. coli* OP50) | 31.43 (compared to *raga-1(ok386)* + *E. coli* OP50) | 41/70 | Fig S1K |
| | *B. infantis* ATCC 15697 | 47.4613 +/- 0.66487 | <0.0001 (compared to *raga-1(ok386)* + *E. coli* OP50) | 28.85 (compared to *raga-1(ok386)* + *E. coli* OP50) | 69/98 | Fig S1K |
| | *B. longum* NCC 2705 | 43.0234 +/- 0.58569 | <0.0001 (compared to *raga-1(ok386)* + *E. coli* OP50) | 16.80 (compared to *raga-1(ok386)* + *E. coli* OP50) | 77/115 | Fig S1K |
| *rict-1(ft7)* | *E. coli* OP50 | 26.2927 +/- 0.84542 | | | 57/78 | |
| | *B. breve* DSMZ 20213 | 37.3413 +/- 0.93508 | <0.0001 (compared to *rict-1(ft7)* + *E. coli* OP50) | 42.02 (compared to *rict-1(ft7)* + *E. coli* OP50) | 73/99 | |
| | *B. infantis* ATCC 15697 | 37.2674 +/- 0.75467 | <0.0001 (compared to *rict-1(ft7)* + *E. coli* OP50) | 41.74 (compared to *rict-1(ft7)* + *E. coli* OP50) | 83/108 | |
| | *B. longum* NCC 2705 | 34.3543 +/- 0.86877 | <0.0001 (compared to *rict-1(ft7)* + *E. coli* OP50) | 30.66 (compared to *rict-1(ft7)* + *E. coli* OP50) | 78/96 | |

| Worm strains | Treatments | Mean +/- s.e.m. (since hatching unless otherwise noted) | *P* values (log-rank) | % lifespan change (compared to the control diet, unless otherwise noted) | # worms (total number of worms died in this assay/(total died + total censored) | Figures |
|---|---|---|---|---|---|---|
| wild-type (N2) | Empty vector + *E. coli* HT115 (control) | 24.2917 +/- 0.60223 | | | 91/101 | Fig 2A, 2G |
| | Empty vector + *B. breve* DSMZ 20213 | 26.8356 +/- 0.83847 | 0.0099 | 10.47 | 77/101 | Fig 2A |
| | Empty vector + *B. infantis* ATCC 15697 | 28.0523 +/- 0.81537 | <0.0001 | 15.48 | 79/95 | Fig 2A, 2G |
| | Empty vector + *B. longum* NCC 2705 | 30.8388 +/- 0.82685 | <0.0001 | 26.95 | 71/102 | Fig 2A |
| | *xbp-1* RNAi + *E. coli* HT115 | 23.0551 +/- 0.4458 | <0.0001 | | 127/129 | Fig 2A, 2G |
| | *xbp-1* RNAi + *B. breve* DSMZ 20213 | 27.0407 +/- 0.91226 | <0.0001 (compared to *xbp-1* RNAi + *E. coli* HT115) | 17.29 (compared to *xbp-1* RNAi + *E. coli* HT115) | 93/112 | Fig 2A |
| | *xbp-1* RNAi + *B. infantis* ATCC 15697 | 23.569 +/- 0.6405 | 0.2523 (compared to *xbp-1* RNAi + *E. coli* HT115) | 2.23 (compared to *xbp-1* RNAi + *E. coli* HT115) | 103/114 | Fig 2A, 2G |
| | *xbp-1* RNAi + *B. longum* NCC 2705 | 29.0846 +/- 1.10659 | <0.0001 (compared to *xbp-1* RNAi + *E. coli* HT115) | 26.15 (compared to *xbp-1* RNAi + *E. coli* HT115) | 43/60 | Fig 2A |
| wild-type (N2) | Empty vector + *E. coli* HT115 (control) | 23.4901 +/- 0.7475 | | | 54/64 | Fig 2A, 2G, 3G, 3I |
| | Empty vector + *B. breve* DSMZ 20213 | 27.6641 +/- 1.04906 | 0.0008 | 17.77 | 58/74 | Fig 2A |
| | Empty vector + *B. infantis* ATCC 15697 | 31.0229 +/- 1.30876 | <0.0001 | 32.07 | 28/32 | Fig 2A, 3G, 3I |
| | Empty vector + *B. longum* NCC 2705 | 29.8171 +/- 1.21578 | <0.0001 | 26.93 | 37/53 | Fig 2A, 2G |
| | *xbp-1* RNAi + *E. coli* HT115 | 23.5883 +/- 0.70295 | | | 76/83 | Fig 2A, 2G |
| | *xbp-1* RNAi + *B. breve* DSMZ 20213 | 28.5962 +/- 1.74736 | 0.0008 (compared to *xbp-1* RNAi + *E. coli* HT115) | 21.23 (compared to *xbp-1* RNAi + *E. coli* HT115) | 30/38 | Fig 2A |
| | *xbp-1* RNAi + *B. infantis* ATCC 15697 | 25.6688 +/- 1.45952 | 0.069 (compared to *xbp-1* RNAi + *E. coli* HT115) | 8.82 (compared to *xbp-1* RNAi + *E. coli* HT115) | 32/43 | Fig 2A |
| | *xbp-1* RNAi + *B. longum* NCC 2705 | 29.3624 +/- 0.92728 | <0.0001 (compared to *xbp-1* RNAi + *E. coli* HT115) | 24.48 (compared to *xbp-1* RNAi + *E. coli* HT115) | 33/42 | Fig 2A, 2G |
| | *fat-7* RNAi + *E. coli* HT115 | 23.5423 +/- 0.72887 | | | 80/84 | Fig 3G, 3I |
| | *fat-7* RNAi + *B. infantis* ATCC 15697 | 25.7942 +/- 0.93856 | 0.0399 (compared to *fat-7* RNAi + *E. coli* HT115) | 9.57 (compared to *fat-7* RNAi + *E. coli* HT115) | 62/67 | Fig 3G, 3I |
| | *fat-7* RNAi + *B. longum* NCC 2705 | 27.4375 +/- 1.36557 | 0.0111 (compared to *fat-7* RNAi + *E. coli* HT115) | 16.55 (compared to *fat-7* RNAi + *E. coli* HT115) | 32/32 | |
| | *gst-4* RNAi + *E. coli* HT115 | 21.6808 +/- 0.57668 | | | 73/79 | Fig 3G |
| | *gst-4* RNAi + *B. infantis* ATCC 15697 | 30.025 +/- 1.50326 | <0.0001 (compared to *gst-4* RNAi + *E. coli* HT115) | 38.49 (compared to *gst-4* RNAi + *E. coli* HT115) | 34/46 | Fig 3G |
| wild-type (N2) | *E. coli* OP50 (control) | 25.7664 +/- 0.35298 | | | 150/165 | Fig 2A, S1G, S1H, S1I |
| | *B. breve* DSMZ 20213 | 28.8616 +/- 0.86026 | <0.0001 | 12.01 | 54/66 | Fig 2A, S1G |
| | *B. infantis* ATCC 15697 | 29.0852 +/- 0.66559 | <0.0001 | 12.88 | 83/112 | Fig S1H |
| | *B. longum* NCC 2705 | 35.6923 +/- 1.21626 | <0.0001 | 38.52 | 13/22 | Fig 2A, S1I |
| *hsf-1(sy441)* | *E. coli* OP50 | 21.1502 +/- 0.33547 | | | 118/132 | Fig 2A |
| | *B. breve* DSMZ 20213 | 25.1844 +/- 0.43429 | <0.0001 (compared to *hsf-1(sy441)* + *E. coli* OP50) | 19.07 (compared to *hsf-1(sy441)* + *E. coli* OP50) | 96/112 | Fig 2A |
| | *B. longum* NCC 2705 | 24.8479 +/- 0.42638 | <0.0001 (compared to *hsf-1(sy441)* + *E. coli* OP50) | 15.78 (compared to *hsf-1(sy441)* + *E. coli* OP50) | 98/108 | Fig 2A |
| *hlh-30(tm1978)* | *E. coli* OP50 | 22.6393 +/- 0.4239 | | | 94/103 | Fig S1G, S1H, S1I |
| | *B. breve* DSMZ 20213 | 20.1191 +/- 0.57989 | 0.0012 (compared to *hlh-30(tm1978)* + *E. coli* OP50) | -11.13 (compared to *hlh-30(tm1978)* + *E. coli* OP50) | 46/64 | Fig S1G |
| | *B. infantis* ATCC 15697 | 19.8525 +/- 0.64724 | <0.0001 (compared to *hlh-30(tm1978)* + *E. coli* OP50) | -12.31 (compared to *hlh-30(tm1978)* + *E. coli* OP50) | 57/73 | Fig S1H |
| | *B. longum* NCC 2705 | 20.414 +/- 0.57647 | 0.0028 (compared to *hlh-30(tm1978)* + *E. coli* OP50) | -9.83 (compared to *hlh-30(tm1978)* + *E. coli* OP50) | 36/48 | Fig S1I |
| wild-type (N2) | *E. coli* OP50 (control) | 25.1496 +/- 0.46084 | | | 119/125 | Fig 2A, S1J, S1M |
| | *B. breve* DSMZ 20213 | 28.0877 +/- 0.61134 | <0.0001 | 11.68 | 77/83 | Fig S1J, S1M |
| | *B. infantis* ATCC 15697 | 29.649 +/- 0.56342 | <0.0001 | 17.89 | 106/110 | Fig 2A, S1J, S1M |
| | *B. longum* NCC 2705 | 33.9497 +/- 0.57938 | <0.0001 | 34.99 | 106/112 | Fig 2A, S1J, S1M |
| *atfs-1(gk3094)* | *E. coli* OP50 | 22.5275 +/- 0.38257 | | | 109/129 | Fig 2A |
| | *B. infantis* ATCC 15697 | 24.0495 +/- 0.46258 | 0.0147 (compared to *atfs-1(gk3094)* + *E. coli* OP50) | 6.76 (compared to *atfs-1(gk3094)* + *E. coli* OP50) | 99/105 | Fig 2A |
| | *B. longum* NCC 2705 | 27.8879 +/- 0.68802 | <0.0001 (compared to *atfs-1(gk3094)* + *E. coli* OP50) | 23.79 (compared to *atfs-1(gk3094)* + *E. coli* OP50) | 53/61 | Fig 2A |

| Worm strains | Treatments | Mean +/- s.e.m. (since hatching unless otherwise noted) | *P* values (log-rank) | % lifespan change (compared to the control diet, unless otherwise noted) | # worms (total number of worms died in this assay/(total died + total censored) | Figures |
|---|---|---|---|---|---|---|
| *eat-2(ad465)* | *E. coli* OP50 | 26.8828 +/- 0.52061 | | | 79/83 | Fig S1J |
| | *B. breve* DSMZ 20213 | 29.8995 +/- 0.90966 | <0.0001 (compared to *eat-2(ad465)* + *E. coli* OP50) | 11.22 (compared to *eat-2(ad465)* + *E. coli* OP50) | 57/79 | Fig S1J |
| | *B. infantis* ATCC 15697 | 33.6269 +/- 0.56463 | <0.0001 (compared to *eat-2(ad465)* + *E. coli* OP50) | 25.09 (compared to *eat-2(ad465)* + *E. coli* OP50) | 94/116 | Fig S1J |
| | *B. longum* NCC 2705 | 32.7919 +/- 0.78362 | <0.0001 (compared to *eat-2(ad465)* + *E. coli* OP50) | 21.98 (compared to *eat-2(ad465)* + *E. coli* OP50) | 86/99 | Fig S1J |
| *rict-1(ft7)* | *E. coli* OP50 | 29.4312 +/- 0.64113 | | | 140/143 | Fig S1M |
| | *B. breve* DSMZ 20213 | 41.5896 +/- 0.92547 | <0.0001 (compared to *rict-1(ft7)* + *E. coli* OP50) | 41.31 (compared to *rict-1(ft7)* + *E. coli* OP50) | 104/105 | Fig S1M |
| | *B. infantis* ATCC 15697 | 38.8103 +/- 0.7049 | <0.0001 (compared to *rict-1(ft7)* + *E. coli* OP50) | 31.87 (compared to *rict-1(ft7)* + *E. coli* OP50) | 147/148 | Fig S1M |
| | *B. longum* NCC 2705 | 43.0597 +/- 0.81944 | <0.0001 (compared to *rict-1(ft7)* + *E. coli* OP50) | 46.31 (compared to *rict-1(ft7)* + *E. coli* OP50) | 147/160 | Fig S1M |
| wild-type (N2) | *E. coli* OP50 (control) | 22.6194 +/- 0.44325 (adult days) | | | 114/127 | Fig S1N |
| | *B. longum* NCC 2705 | 30.6274 +/- 1.08336 (adult days) | <0.0001 | 35.40 | 61/80 | Fig S1N |
| *nuo-6(qm200)* | *E. coli* OP50 | 33.1088 +/- 0.79261 (adult days) | | | 93/106 | Fig S1N |
| | *B. longum* NCC 2705 | 37.5588 +/- 1.42776 (adult days) | 0.0045 (compared to *nuo-6(qm200)* + *E. coli* OP50) | 13.44 (compared to *nuo-6(qm200)* + *E. coli* OP50) | 36/51 | Fig S1N |
| *isp-1(qm150)* | *E. coli* OP50 | 31.3937+/- 0.94802 (adult days) | | | 103/117 | Fig S1N |
| | *B. longum* NCC 2705 | 33.684 +/- 1.22857 (adult days) | 0.1001 (compared to *isp-1(qm150)* + *E. coli* OP50) | 7.30 (compared to *isp-1(qm150)* + *E. coli* OP50) | 57/92 | Fig S1N |
| wild-type (N2) | *E. coli* OP50 (control) | 20.7614 +/- 0.49407 (adult days) | | | 70/80 | Fig S1N |
| | *B. infantis* ATCC 15697 | 26.1756 +/- 0.7762 (adult days) | <0.0001 | 26.08 | 50/56 | Fig S1N |
| *nuo-6(qm200)* | *E. coli* OP50 | 32.4323 +/- 0.76514 (adult days) | | | 72/80 | Fig S1N |
| | *B. infantis* ATCC 15697 | 37.2575 +/- 0.88727 (adult days) | 0.0002 (compared to *nuo-6(qm200)* + *E. coli* OP50) | 14.88 (compared to *nuo-6(qm200)* + *E. coli* OP50) | 57/61 | Fig S1N |
| *isp-1(qm150)* | *E. coli* OP50 | 29.0093 +/- 1.12871 (adult days) | | | 60/77 | Fig S1N |
| | *B. infantis* ATCC 15697 | 31.9199 +/- 1.06004 (adult days) | 0.2853 (compared to *isp-1(qm150)* + *E. coli* OP50) | 10.03 (compared to *isp-1(qm150)* + *E. coli* OP50) | 47/56 | Fig S1N |
| wild-type (N2) | *E. coli* OP50 (control) | 24.7318 +/- 0.41996 | | | 100/112 | Fig 3G |
| | *B. infantis* ATCC 15697 | 27.7905 +/- 0.52415 | <0.0001 | 12.37 | 99/120 | Fig 3G |
| *sod-4(gk101)* | *E. coli* OP50 | 27.1225 +/- 0.54221 | | | 121/131 | Fig 3G |
| | *B. infantis* ATCC 15697 | 30.9919 +/- 0.78815 | <0.0001 (compared to *sod-4(gk101)* + *E. coli* OP50) | 14.27 (compared to *sod-4(gk101)* + *E. coli* OP50) | 95/112 | Fig 3G |
| wild-type (N2) | Empty vector + *E. coli* HT115 (control) | 19.0492 +/- 0.48972 | | | 92/97 | Fig 2A, 3G |
| | Empty vector + *B. infantis* ATCC 15697 | 24.3575 +/- 0.79489 | <0.0001 | 27.87 | 58/80 | Fig 2A, 3G |
| | Empty vector + *B. longum* NCC 2705 | 24.5221 +/- 0.89302 | <0.0001 | 28.73 | 42/72 | Fig 2A |
| | *sbp-1* RNAi + *E. coli* HT115 (RNAi only on adult day one) | 15.7805 +/- 0.52209 | | | 41/42 | Fig 2A |
| | *sbp-1* RNAi + *B. infantis* ATCC 15697 (RNAi only on adult day one) | 23.2234 +/- 1.08984 | <0.0001 (compared to *sbp-1* RNAi + *E. coli* HT115) | 47.17 (compared to *sbp-1* RNAi + *E. coli* HT115) | 26/29 | Fig 2A |
| | *sbp-1* RNAi + *B. longum* NCC 2705 (RNAi only on adult day one) | 23.2908 +/- 1.28848 | <0.0001 (compared to *sbp-1* RNAi + *E. coli* HT115) | 47.59 (compared to *sbp-1* RNAi + *E. coli* HT115) | 26/32 | Fig 2A |
| | *lipl-3* RNAi + *E. coli* HT115 | 19.4207 +/- 0.54065 | | | 72/74 | Fig 3G |
| | *lipl-3* RNAi + *B. infantis* ATCC 15697 | 23.9593 +/- 0.70909 | <0.0001 (compared to *lipl-3* RNAi + *E. coli* HT115) | 23.37 (compared to *lipl-3* RNAi + *E. coli* HT115) | 57/63 | Fig 3G |
| wild-type (N2) | *E. coli* OP50 (control) | 23.2291 +/- 0.27201 | | | 119/121 | Fig 2A, S1E, S2D |
| | *B. breve* DSMZ 20213 | 28.001 +/- 0.595367 | <0.0001 | 20.54 | 87/105 | Fig 2A, S1E |
| | *B. infantis* ATCC 15697 | 27.3207 +/- 0.77521 | <0.0001 | 17.61 | 56/60 | Fig S2D |
| | *B. longum* NCC 2705 | 27.8202 +/- 0.64476 | <0.0001 | 19.76 | 74/81 | |

| Worm strains | Treatments | Mean +/- s.e.m. (since hatching unless otherwise noted) | *P* values (log-rank) | % lifespan change (compared to the control diet, unless otherwise noted) | # worms (total number of worms died in this assay/(total died + total censored) | Figures |
|---|---|---|---|---|---|---|
| *nhr-49(nr2041)* | *E. coli* OP50 | 17.8275 +/- 0.2919 | | | 127/128 | Fig 2A |
| | *B. breve* DSMZ 20213 | 16.3024 +/- 0.32658 | 0.0014 (compared to *nhr-49(nr2041)* + *E. coli* OP50) | -5.48 (compared to *nhr-49(nr2041)* + *E. coli* OP50) | 106/108 | Fig 2A |
| *daf-16(mu86)* | *E. coli* OP50 | 20.3599 +/- 0.28746 | | | 94/100 | Fig 2A, S1E |
| | *B. breve* DSMZ 20213 | 20.2298 +/- 0.36359 | 0.7548 (compared to *daf-16(mu86)* + *E. coli* OP50) | -0.64 (compared to *daf-16(mu86)* + *E. coli* OP50) | 87/100 | Fig 2A, S1E |
| *fat-6(tm331);fat-7(wa37)* | *E. coli* OP50 | 23.4199 +/- 0.71572 | | | 52/84 | Fig S2D |
| | *B. infantis* ATCC 15697 | 24.2385 +/- 0.65385 | 0.5622 (compared to *fat-6(tm331);fat-7(wa37)* + *E. coli* OP50) | 3.50 (compared to *fat-6(tm331);fat-7(wa37)* + *E. coli* OP50) | 60/70 | Fig S2D |
| wild-type (N2) | *E. coli* OP50 (control) | 24.6194 +/- 0.44325 | | | 114/127 | Fig S2G |
| | *B. longum* NCC 2705 | 32.6274 +/- 1.08336 | <0.0001 | 32.53 | 61/80 | Fig S2G |
| *fat-6(tm331);fat-7(wa37)* | *E. coli* OP50 | 26.015 +/- 0.64611 | | | 50/71 | Fig S2G |
| | *B. longum* NCC 2705 | 27.2889 +/- 1.0223 | 0.3066 (compared to *fat-6(tm331);fat-7(wa37)* + *E. coli* OP50) | 4.90 (compared to *fat-6(tm331);fat-7(wa37)* + *E. coli* OP50) | 42/59 | Fig S2G |
| wild-type (N2) | Empty vector + *E. coli* HT115 (control) | 20.3724 +/- 0.59315 | | | 70/75 | Fig 2A, S2H |
| | Empty vector + *B. infantis* ATCC 15697 | 22.8414 +/- 1.20431 | 0.0175 | 12.12 | 22/47 | Fig 2A |
| | Empty vector + *B. longum* NCC 2705 | 27.4224 +/- 1.06787 | <0.0001 | 34.61 | 38/57 | Fig 2A, S2H |
| | *sbp-1* RNAi + *E. coli* HT115 (RNAi only on adult day one) | 16.2614 +/- 0.33423 | | | 72/74 | Fig 2A |
| | *sbp-1* RNAi + *B. infantis* ATCC 15697 (RNAi only on adult day one) | 22.2025 +/- 0.72954 | <0.0001 (compared to *sbp-1* RNAi + *E. coil* HT115) | 36.53 (compared to *sbp-1* RNAi + *E. coil* HT115) | 46/57 | Fig 2A |
| | *sbp-1* RNAi + *B. longum* NCC 2705 (RNAi only on adult day one) | 21.4718 +/- 0.82377 | <0.0001 (compared to *sbp-1* RNAi + *E. coil* HT115) | 32.04 (compared to *sbp-1* RNAi + *E. coil* HT115) | 47/57 | Fig 2A |
| | *fat-2* RNAi + *E. coli* HT115 (RNAi only on adult day one) | 18.884 +/- 0.53223 | | | 70/75 | |
| | *fat-2* RNAi + *B. infantis* ATCC 15697 (RNAi only on adult day one) | 23.1778 +/- 1.21346 | 0.0017 (compared to *fat-2* RNAi + *E. coli* HT115) | 22.74 (compared to *fat-2* RNAi + *E. coli* HT115) | 17/18 | |
| | *fat-7* RNAi + *E. coli* HT115 | 21.8569 +/- 0.71994 | | | 63/71 | Fig S2H |
| | *fat-7* RNAi + *B. infantis* ATCC 15697 | 20.8026 +/- 0.95055 | 0.2573 (compared to *fat-7* RNAi + *E. coil* HT115) | -4.82 (compared to *fat-7* RNAi + *E. coil* HT115) | 27/57 | |
| | *fat-7* RNAi + *B. longum* NCC 2705 | 22.7838 +/- 1.11128 | 0.5643 (compared to *fat-7* RNAi + *E. coil* HT115) | 4.24 (compared to *fat-7* RNAi + *E. coil* HT115) | 24/62 | Fig S2H |
| wild-type (N2) | *E. coli* OP50 (control) | 23.0574 +/- 0.50643 | | | 98/114 | Fig 2A |
| | *B. infantis* ATCC 15697 | 29.673 +/- 0.64473 | <0.0001 | 28.69 | 106/128 | Fig 2A |
| | *B. longum* NCC 2705 | 30.0149 +/- 0.68457 | <0.0001 | 30.17 | 83/116 | Fig 2A |
| *daf-16(mu86)* | *E. coli* OP50 | 19.3896 +/- 0.31265 | | | 103/117 | Fig 2A |
| | *B. infantis* ATCC 15697 | 20.1816 +/- 0.66298 | 0.2052 (compared to *daf-16(mu86)* + *E. coli* OP50) | 4.08 (compared to *daf-16(mu86)* + *E. coli* OP50) | 53/89 | Fig 2A |
| | *B. longum* NCC 2705 | 24.1685 +/- 0.86352 | <0.0001 (compared to *daf-16(mu86)* + *E. coli* OP50) | 24.65 (compared to *daf-16(mu86)* + *E. coli* OP50) | 47/77 | Fig 2A |
| wild-type (N2) | *E. coli* OP50 (control) | 23.0037 +/- 0.45602 | | | 104/106 | Fig 2A |
| | *B. breve* DSMZ 20213 | 25.8722 +/- 0.55492 | <0.0001 | 12.47 | 67/88 | Fig 2A |
| | *B. infantis* ATCC 15697 | 26.73 +/- 0.5932 | <0.0001 | 16.20 | 82/109 | Fig 2A |
| | *B. longum* NCC 2705 | 30.2691 +/- 0.68895 | <0.0001 | 31.58 | 32/47 | Fig 2A |
| *nhr-80(tm1011)* | *E. coli* OP50 | 22.8295 +/- 0.45941 | | | 58/62 | Fig 2A |
| | *B. breve* DSMZ 20213 | 29.1459 +/- 0.69024 | <0.0001 (compared to *nhr-80(tm1011)* + *E. coli* OP50) | 27.67 (compared to *nhr-80(tm1011)* + *E. coli* OP50) | 56/77 | Fig 2A |
| | *B. infantis* ATCC 15697 | 29.8814 +/- 0.87612 | <0.0001 (compared to *nhr-80(tm1011)* + *E. coli* OP50) | 30.89 (compared to *nhr-80(tm1011)* + *E. coli* OP50) | 61/70 | Fig 2A |
| | *B. longum* NCC 2705 | 28.6967 +/- 0.62133 | <0.0001 (compared to *nhr-80(tm1011)* + *E. coli* OP50) | 25.70 (compared to *nhr-80(tm1011)* + *E. coli* OP50) | 56/70 | Fig 2A |

| Worm strains | Treatments | Mean +/- s.e.m.<br>(since hatching unless otherwise noted) | *P* values<br>(log-rank) | % lifespan change<br>(compared to the control diet, unless otherwise noted) | # worms<br>(total number of worms died in this assay/(total died + total censored) | Figures |
|---|---|---|---|---|---|---|
| wild-type (N2) | Empty vector + *E. coli* HT115 (control) | 26.6879 +/- 1.51945 | | | 21/29 | Fig 2A, 3G |
| | Empty vector + *B. breve* DSMZ 20213 | 34.3056 +/- 1.20638 | 0.0001 | 28.54 | 32/43 | Fig 2A |
| | Empty vector + *B. infantis* ATCC 15697 | 32.3755 +/- 1.03656 | 0.0017 | 21.31 | 43/52 | Fig 3G |
| | Empty vector + *B. longum* NCC 2705 | 32.0119 +/- 1.51034 | 0.016 | 19.95 | 33/45 | Fig 2A |
| | *sbp-1* RNAi + *E. coli* HT115<br>(RNAi only on adult day one) | 22.7584 +/- 1.21178 | | | 23/26 | Fig 2A |
| | *sbp-1* RNAi + *B. breve* DSMZ 20213<br>(RNAi only on adult day one) | 30.5984 +/- 1.19106 | <0.0001<br>(compared to *sbp-1* RNAi + *E. coil* HT115) | 34.45<br>(compared to *sbp-1* RNAi + *E. coil* HT115) | 32/44 | Fig 2A |
| | *sbp-1* RNAi + *B. longum* NCC 2705<br>(RNAi only on adult day one) | 27.8991 +/- 0.98517 | 0.0011<br>(compared to *sbp-1* RNAi + *E. coil* HT115) | 22.59<br>(compared to *sbp-1* RNAi + *E. coil* HT115) | 43/50 | Fig 2A |
| | *fat-2* RNAi + *E. coli* HT115<br>(RNAi only on adult day one) | 24.071 +/- 1.09032 | | | 41/44 | Fig 3G |
| | *fat-2* RNAi + *B. infantis* ATCC 15697<br>(RNAi only on adult day one) | 29.7083 +/- 1.40772 | 0.0027<br>(compared to *fat-2* RNAi + *E. coil* HT115) | 23.42<br>(compared to *fat-2* RNAi + *E. coil* HT115) | 29/31 | Fig 3G |
| wild-type (N2) | *E. coli* OP50 (control) | 25.2707 +/- 0.4404 | | | 87/96 | |
| | *B. breve* DSMZ 20213 | 27.0456 +/- 0.68846 | 0.004 | 7.02 | 76/97 | |
| | *B. infantis* ATCC 15697 | 30.848 +/- 0.95617 | <0.0001 | 23.97 | 57/70 | |
| | *B. longum* NCC 2705 | 34.2578 +/- 0.65198 | <0.0001 | 38.62 | 83/92 | |
| *hlh-30(tm1978)* | *E. coli* OP50 | 19.7502 +/- 0.41396 | | | 117/130 | |
| | *B. breve* DSMZ 20213 | 17.3309 +/- 0.45736 | 0.0002<br>(compared to *hlh-30(tm1978)* + *E. coli* OP50) | -12.25<br>(compared to *hlh-30(tm1978)* + *E. coli* OP50) | 84/86 | |
| | *B. infantis* ATCC 15697 | 17.1179 +/- 0.42734 | <0.0001<br>(compared to *hlh-30(tm1978)* + *E. coli* OP50) | -13.33<br>(compared to *hlh-30(tm1978)* + *E. coli* OP50) | 115/123 | |
| | *B. longum* NCC 2705 | 18.4885 +/- 0.48437 | 0.0833<br>(compared to *hlh-30(tm1978)* + *E. coli* OP50) | -6.39<br>(compared to *hlh-30(tm1978)* + *E. coli* OP50) | 94/103 | |
| wild-type (N2) | *E. coli* OP50 (control) | 23.9098 +/- 0.42996 | | | 114/121 | |
| | *B. breve* DSMZ 20213 | 27.4561 +/- 0.61431 | <0.0001 | 14.83 | 83/106 | |
| | *B. infantis* ATCC 15697 | 31.2595 +/- 0.93603 | <0.0001 | 30.74 | 69/85 | |
| | *B. longum* NCC 2705 | 35.9605 +/- 1.41245 | <0.0001 | 50.40 | 21/24 | |
| *eat-2(ad465)* | *E. coli* OP50 | 25.8349 +/- 0.46901 | | | 96/101 | |
| | *B. infantis* ATCC 15697 | 35.3455 +/- 0.85395 | <0.0001<br>(compared to *eat-2(ad465)* + *E. coli* OP50) | 36.81<br>(compared to *eat-2(ad465)* + *E. coli* OP50) | 61/86 | |
| wild-type (N2) | *E. coli* OP50 (control) | 25.43 +/- 0.76191 (adult days) | | | 54/64 | |
| | *B. longum* NCC 2705 | 32.4862 +/- 1.42802 (adult days) | <0.0001 | 27.75 | 39/42 | |
| *nuo-6(qm200)* | *E. coli* OP50 | 31.9119 +/- 0.80746 (adult days) | | | 107/121 | |
| | *B. longum* NCC 2705 | 36.5426 +/- 0.84355 (adult days) | 0.0012<br>(compared to *nuo-6(qm200)* + *E. coli* OP50) | 14.51<br>(compared to *nuo-6(qm200)* + *E. coli* OP50) | 73/78 | |
| wild-type (N2) | *E. coli* OP50 (control) | 23.2707 +/- 0.4404 (adult days) | | | 87/96 | |
| | *B. infantis* ATCC 15697 | 28.848 +/- 0.95617 (adult days) | <0.0001 | 23.97 | 57/70 | |
| | *B. longum* NCC 2705 | 32.2578 +/- 0.65198 (adult days) | <0.0001 | 38.62 | 83/92 | |
| *isp-1(qm150)* | *E. coli* OP50 | 30.1975 +/- 1.08968 (adult days) | | | 66/85 | |
| | *B. infantis* ATCC 15697 | 33.1759 +/- 1.28516 (adult days) | 0.2602<br>(compared to *isp-1(qm150)* + *E. coli* OP50) | 9.86<br>(compared to *isp-1(qm150)* + *E. coli* OP50) | 35/48 | |
| | *B. longum* NCC 2705 | 31.7924 +/- 1.42384 (adult days) | 0.2941<br>(compared to *isp-1(qm150)* + *E. coli* OP50) | 5.28<br>(compared to *isp-1(qm150)* + *E. coli* OP50) | 44/54 | |
| wild-type (N2) | Empty vector + *E. coli* HT115 (control) | 24.2932 +/- 0.56645 | | | 77/95 | |
| | Empty vector + *B. infantis* ATCC 15697 | 29.7364 +/- 0.63426 | <0.0001 | 22.41 | 80/93 | |
| | *fat-7* RNAi + *E. coli* HT115 | 22.0981 +/- 0.52851 | | | 62/72 | |
| | *fat-7* RNAi + *B. infantis* ATCC 15697 | 24.8991 +/- 0.59528 | 0.0006<br>(compared to *fat-7* RNAi + *E. coli* HT115) | 12.68<br>(compared to *fat-7* RNAi + *E. coli* HT115) | 75/85 | |

**Additional information:** Experiments corresponding to each figure are indicated in the right-most column. Mean lifespan and s.e.m. were calculated from independent replicate plates. *P* values were determined by log-rank test using pooled worms from independent replicate plates. "# worms" indicates total deaths/(total deaths + total censored). Censored worms included animals that underwent internal hatching ("bagging"), exhibited a ruptured vulva, or crawled off the plates.

**S4 Table: Summary of oxidative stress survival statistics**

| Worm strains | Treatments | Mean +/- s.e.m. (hours) | *P* values (log-rank) | % survival change (compared to the control unless otherwise noted) | # worms (total number of worms died in this assay/(total died + total censored) | Figures |
|---|---|---|---|---|---|---|
| wild-type (N2) | *E. coli* OP50 (control) | 6.70492 +/- 0.22096 | | | 61/61 | Fig 1I |
| | *B. longum* NCC 2705 | 10.2222 +/- 0.29634 | <0.0001 | 52.46 | 38/54 | Fig 1I |
| | *B. infantis* ATCC 15697 | 8.97297 +/- 0.50708 | <0.0001 | 33.83 | 27/37 | Fig 1I |
| | *B. breve* DSMZ 20213 | 7.58621 +/- 0.29825 | 0.0031 | 13.14 | 58/58 | Fig 1I |
| wild-type (N2) | *E. coli* OP50 (control) | 7.21429 +/- 0.39809 | | | 52/56 | Fig S1C |
| | *B. adolescentis* L2-32 | 4.36508 +/- 0.32893 | <0.0001 | -39.49 | 63/63 | Fig S1C |
| | *B. angulatum* DSMZ 20098 | 5.63265 +/- 0.46456 | 0.0246 | -21.92 | 48/49 | Fig S1C |
| | *B. infantis* ATCC 15697 | 8.61667 +/- 0.3678 | 0.0044 | 19.44 | 45/60 | Fig S1C |
| | *B. longum* NCC 2705 | 9.12281 +/- 0.35774 | 0.0005 | 26.45 | 43/57 | Fig S1C |
| wild-type (N2) | *E. coli* OP50 (control) | 6.25676 +/- 0.29563 | | | 74/74 | Fig 3A, B, C |
| | *B. infantis* ATCC 15697 | 7.88312 +/- 0.3996 | <0.0001 | 25.99 | 64/80 | Fig 3A, B, C |
| *atfs-1(gk3094)* | *E. coli* OP50 | 4.82353 +/- 0.28657 | | | 51/51 | Fig 3B |
| | *B. infantis* ATCC 15697 | 5.72549 +/- 0.37797 | 0.0625 (compared to *atfs-1(gk3094)* + *E. coli* OP50) | 18.70 (compared to *atfs-1(gk3094)* + *E. coli* OP50) | 50/51 | Fig 3B |
| *skn-1(mg570)* | *E. coli* OP50 | 6.98765 +/- 0.31852 | | | 77/81 | Fig 3C |
| | *B. infantis* ATCC 15697 | 7.49333 +/- 0.35487 | 0.0694 (compared to *skn-1(mg570)* + *E. coli* OP50) | 7.24 (compared to *skn-1(mg570)* + *E. coli* OP50) | 61/75 | Fig 3C |
| *nhr-49(nr2041)* | *E. coli* OP50 | 3.61017 +/- 0.30867 | | | 59/59 | Fig 3A |
| | *B. infantis* ATCC 15697 | 3.33962 +/- 0.36536 | 0.7709 (compared to *nhr-49(nr2041)* + *E. coli* OP50) | -7.49 (compared to *nhr-49(nr2041)* + *E. coli* OP50) | 53/53 | Fig 3A |
| wild-type (N2) | Empty vector + *E. coli* HT115 (control) | 7.08929 +/- 0.4687 | | | 48/56 | Fig 3D, 3J, S2I |
| | Empty vector + *B. infantis* ATCC 15697 | 10.1905 +/- 0.32996 | <0.0001 | 43.75 | 30/63 | Fig 3D, 3J |
| | Empty vector + *B. longum* NCC 2705 | 10.0938 +/- 0.28791 | <0.0001 | 42.38 | 42/64 | Fig S2I |
| | *fat-7* RNAi + *E. coli* HT115 | 7.55357 +/- 0.46166 | | | 46/56 | Fig 3J, S2I |
| | *fat-7* RNAi + *B. infantis* ATCC 15697 | 8.28889 +/- 0.48966 | 0.3129 (compared to *fat-7* RNAi + *E. coli* HT115) | 9.73 (compared to *fat-7* RNAi + *E. coli* HT115) | 34/45 | Fig 3J |
| | *fat-7* RNAi + *B. longum* NCC 2705 | 8.60811 +/- 0.2933 | 0.029 (compared to *fat-7* RNAi + *E. coli* HT115) | 13.96 (compared to *fat-7* RNAi + *E. coli* HT115) | 50/74 | Fig S2I |
| | *xbp-1* RNAi + *E. coli* HT115 | 8.9322 +/- 0.37259 | | | 42/59 | Fig 3D |
| | *xbp-1* RNAi + *B. infantis* ATCC 15697 | 8.57143 +/- 0.33407 | 0.3301 (compared to *xbp-1* RNAi + *E. coli* HT115) | -4.04 (compared to *xbp-1* RNAi + *E. coli* HT115) | 61/77 | Fig 3D |
| wild-type (N2) | *E. coli* OP50 (control) | 6.90769 +/- 0.31678 | | | 65/65 | Fig S2A |
| | *B. longum* NCC 2705 | 9.80303 +/- 0.26985 | <0.0001 | 41.91 | 47/66 | |
| | *B. infantis* ATCC 15697 | 9.07813 +/- 0.3633 | <0.0001 | 31.42 | 45/64 | Fig S2A |
| | *B. breve* DSMZ 20213 | 8 +/- 0.48812 | 0.0051 | 15.81 | 34/40 | |
| *daf-16(mu86)* | *E. coli* OP50 | 5.24675 +/- 0.2883 | | | 76/77 | Fig S2A |
| | *B. infantis* ATCC 15697 | 8.21429 +/- 0.36206 | <0.0001 (compared to *daf-16(mu86)* + *E. coli* OP50) | 56.56 (compared to *daf-16(mu86)* + *E. coli* OP50) | 61/70 | Fig S2A |

| Worm strains | Treatments | Mean +/- s.e.m. (hours) | *P* values (log-rank) | % survival change (compared to the control unless otherwise noted) | # worms (total number of worms died in this assay/(total died + total censored) | Figures |
|---|---|---|---|---|---|---|
| wild-type (N2) | *E. coli* OP50 (control) | 5.90769 +/- 0.27008 | | | 65/65 | Fig S2B |
| | *B. infantis* ATCC 15697 | 7.39706 +/- 0.43567 | <0.0001 | 25.21 | 57/68 | Fig S2B |
| *skn-1(mg570)* | *E. coli* OP50 | 7.31373 +/- 0.42174 | | | 48/51 | |
| | *B. infantis* ATCC 15697 | 7.69231 +/- 0.44955 | 0.4468 (compared to *skn-1(mg570)* + *E. coli* OP50) | 5.18 (compared to *skn-1(mg570)* + *E. coli* OP50) | 48/52 | |
| *hsf-1(sy441)* | *E. coli* OP50 | 4.2459 +/- 0.26308 | | | 61/61 | Fig S2B |
| | *B. infantis* ATCC 15697 | 5.49153 +/- 0.34322 | 0.0023 (compared to *hsf-1(sy441)* + *E. coli* OP50) | 29.42 (compared to *hsf-1(sy441)* + *E. coli* OP50) | 56/59 | Fig S2B |
| *daf-16(mu86)* | *E. coli* OP50 | 5.09804 +/- 0.36435 | | | 51/51 | |
| | *B. infantis* ATCC 15697 | 7.56 +/- 0.44787 | <0.0001 (compared to *daf-16(mu86)* + *E. coli* OP50) | 48.29 (compared to *daf-16(mu86)* + *E. coli* OP50) | 45/50 | |
| *nhr-49(nr2041)* | *E. coli* OP50 | 3.89583 +/- 0.33586 | | | 48/48 | |
| | *B. infantis* ATCC 15697 | 3.06383 +/- 0.24785 | 0.0217 (compared to *nhr-49(nr2041)* + *E. coli* OP50) | -21.36 (compared to *nhr-49(nr2041)* + *E. coli* OP50) | 47/47 | |
| wild-type (N2) | *E. coli* OP50 + 50% MeOH (control) | 7.42308 +/- 0.45841 | | | 42/52 | Fig 4F |
| | *E. coli* OP50 + *B. infantis* ATCC 15697 lipid extract | 9.65909 +/- 0.40648 | 0.013 | 30.12 | 32/44 | Fig 4F |
| | *E. coli* OP50 + *B. longum* NCC 2705 lipid extract | 9.22222 +/- 0.44047 | 0.0158 | 24.24 | 30/45 | Fig 4F |
| wild-type (N2) | *E. coli* OP50 (control) | 6.77358 +/- 0.37597 | | | 51/53 | |
| | *B. infantis* ATCC 15697 | 9.12821 +/- 0.56028 | <0.0001 | 34.76 | 25/39 | |
| *atfs-1(gk3094)* | *E. coli* OP50 | 7.66667 +/- 0.41741 | | | 40/45 | |
| | *B. infantis* ATCC 15697 | 8.44737 +/- 0.57564 | 0.0701 (compared to *atfs-1(gk3094)* + *E. coli* OP50) | 10.18 (compared to *atfs-1(gk3094)* + *E. coli* OP50) | 28/38 | |
| wild-type (N2) | *E. coli* OP50 (control) | 7.73529 +/- 0.39899 | | | 34/34 | |
| | *B. infantis* ATCC 15697 | 8.65854 +/- 0.49139 | 0.0063 | 11.94 | 32/41 | |
| *hsf-1(sy441)* | *E. coli* OP50 | 5.03922 +/- 0.37255 | | | 51/51 | |
| | *B. infantis* ATCC 15697 | 6.33333 +/- 0.42443 | 0.0111 (compared to *hsf-1(sy441)* + *E. coli* OP50) | 25.68 (compared to *hsf-1(sy441)* + *E. coli* OP50) | 58/60 | |
| wild-type (N2) | Empty vector + *E. coli* HT115 (control) | 8.08333 +/- 0.44527 | | | 41/48 | |
| | Empty vector + *B. infantis* ATCC 15697 | 10.16 +/- 0.35296 | 0.0003 | 25.69 | 29/50 | |
| | *xbp-1* RNAi + *E. coli* HT115 | 9.375 +/- 0.38693 | | | 56/56 | |
| | *xbp-1* RNAi + *B. infantis* ATCC 15697 | 9.72727 +/- 0.34078 | 0.6984 (compared to *xbp-1* RNAi + *E. coli* HT115) | 3.76 (compared to *xbp-1* RNAi + *E. coli* HT115) | 55/55 | |
| wild-type (N2) | Empty vecotr + *E. coli* HT115 (control) | 7.57143 +/- 0.44395 | | | 50/56 | |
| | Empty vecotr + *B. longum* NCC 2705 | 10.2333 +/- 0.32057 | <0.0001 | 35.11 | 33/60 | |
| | *fat-7* RNAi + *E. coli* HT115 | 7.93333 +/- 0.47054 | | | 52/60 | |
| | *fat-7* RNAi + *B. longum* NCC 2705 | 10.0702 +/- 0.27385 | 0.0064 (compared to *fat-7* RNAi + *E. coli* HT115) | 26.94 (compared to *fat-7* RNAi + *E. coli* HT115) | 41/57 | |
| wild-type (N2) | *E. coli* OP50 + 50% MeOH (control) | 7.47458 +/- 0.35169 | | | 55/59 | |
| | *E. coli* OP50 + *B. infantis* ATCC 15697 lipid extract | 8.70423 +/- 0.29999 | 0.0322 | 16.45 | 66/71 | |
| wild-type (N2) | *E. coli* OP50 + 50% MeOH (control) | 7.22581 +/- 0.35592 | | | 60/62 | |
| | *E. coli* OP50 + *B. longum* NCC 2705 lipid extract | 8.26027 +/- 0.35157 | 0.0075 | 14.32 | 62/73 | |

**Additional information**: Experiments corresponding to each figure are indicated in the right-most column. Survival was scored hourly for up to 12 hours. Worms that remained alive at the end of the experiment were censored. Mean survival time and s.e.m. were calculated from independent replicate plates. *P* values were determined by log-rank test using pooled worms from independent replicate plates. “# worms” indicates total deaths/(total deaths + total censored). Censored worms were defined as animals still alive at the 12-hour experimental endpoint.

**S5 Table: List of *C. elegans* strains used in this study**

| Strain name | Genotype | Nomenclature used in this study | Original source |
|---|---|---|---|
| N2 | wild-type | N2 | CGC |
| CF1038 | *daf-16(mu86) I* | *daf-16(mu86)* | CGC |
| GR2245 | *skn-1(mg570) IV* | *skn-1(mg570)* | CGC |
| VC3201 | *atfs-1(gk3094) V* | *atfs-1(gk3094)* | CGC |
| PS3551 | *hsf-1(sy441) I* | *hsf-1(sy441)* | CGC |
| STE68 | *nhr-49(nr2041) I* | *nhr-49(nr2041)* | CGC |
| JIN1375 | *hlh-30(tm1978) IV* | *hlh-30(tm1978)* | CGC |
| MQ1333 | *nuo-6(qm200) I* | *nuo-6(qm200)* | CGC |
| MQ887 | *isp-1(qm150) IV* | *isp-1(qm150)* | CGC |
| STE70 | *nhr-80(tm1011) III* | *nhr-80(tm1011)* | CGC |
| KQ1366 | *rict-1(ft7) II* | *rict-1(ft7)* | CGC |
| VC222 | *raga-1(ok386) II* | *raga-1(ok386)* | CGC |
| DAF465 | *eat-2(ad465) II* | *eat-2(ad465)* | CGC |
| RB1206 | *rsks-1(ok1255) III* | *rsks-1(ok1255)* | CGC |
| BX107 | *fat-5(tm420) V* | *fat-5(tm420)* | CGC |
| BX156 | *fat-6(tm331) IV;fat-7(wa36) V* | *fat-6(tm331);fat-7(wa37)* | CGC |
| BX153 | *fat-7(wa36) V* | *fat-7(wa36)* | CGC |
| GA416 | *sod-4(gk101) III* | *sod-4(gk101)* | CGC |
| SJ4005 | *zcIs4 [hsp-4::GFP] V* | *hsp-4p*::GFP | CGC |
| SJ4100 | *zcIs13 [hsp-6p::GFP + lin-15(+)]* | *hsp-6p*::GFP | CGC |

**S6 Table: List of RT-qPCR primers used in this study**

| Gene | Forward primer (5′–3′) | Reverse primer (5′–3′) |
|---|---|---|
| *act-1* | CTCTTGCCCCATCAACCATG | CTTGCTTGGAGATCCACATC |
| *sod-1* | ATCTGGATCACACAGAAGTCCG | GCATCCGTTGGTGGAATCAC |
| *sod-2* | TCACCGCAATTAAGAGCGACT | GCGACAGTTGATGCCGAAAG |
| *sod-3* | ATGGACACTATTAAGCGCGAC | CCAGAGCCTTGAACCGCAAT |
| *sod-4* | TGGCTCTCTCCGTTTGCATT | ACCGATCCGTTAAGCTTCAGA |
| *sod-5* | AACGTGCTGTAGCGGTTCTC | CACCTTCGGCTTTCTGGGTAA |
| *ctl-1* | CAAGGAGACGTATCCAAAACCC | ACGGCGGTCTTCGAGTAGAT |
| *ctl-2* | TCCCAGATGGGTACCGTCAT | TCACTCCTTGAGTTGGCTTGAA |
| *ctl-3* | ATAGAGATAGAAATCAGGAACCCCA | CCACCACCTTTGGCATGGAC |
| *trx-2* | TTTCTCACGGCGCTTCTGTT | CCTTGTCTTCCGTTTACCTTTTCC |
| *prdx-2* | ACCAACCACCAAATCTCCCG | CGATGATGAAGAGTCCACGGAA |
| *prdx-3* | TGATAAGCACGGAGAGGTTTGC | GGTAGCGACTCCTGGCTTG |
| *gst-4* | CTCTTGCTGAGCCAATCCGT | CCAAATGGAGTCGTTGGCTTC |
| *gst-8* | CCGTGGAGCTGGAGAGGTTA | CGAATGGGGTGGTTGGTTTG |
| *gst-10* | ACTACTTCACTATTCGAGGATTCGG | GCATGGCAACTGACCAAGGA |
| *gcs-1* | GCAGGTGAATGCGATGCTTG | CAAGCGATGAGACCTCCGT |
| *fmo-2* | TGTCAAAATGGGGAACAAGCG | CCATAGAGAAGACCATGTCGAATC |
| *fat-1* | ATGTCGCTGAGGTGTACGAG | TCCGTCTGTGATATGGTGCA |
| *fat-2* | AGGATATTGAGGTCTACGAAGCT | TGCATGATGTTGTCGAGTCC |
| *fat-3* | TATCAGATCGAGCACCACTTGT | TCATCGACGAGGTAAGGAAGA |
| *fat-4* | ACTATCAGATTGAGCACCATCTT | CCTGTGAAATAATCGTCGACC |
| *fat-5* | GCGATTTGTACGAGGATCCG | AGGACGACTGGAATGAAGGT |
| *fat-6* | CGCTGCTCACTATTTCGGAT | AAGTTGTGACCTCCCTCTCC |
| *fat-7* | GCGCTGCTCACTATTTTGGTT | CCTCCTTCACCAACGGCTAC |
| *elo-1* | GCACTTCACCAATGCCAACT | CCGCGGAGAACATATGATTGG |
| *elo-2* | TTGCTCTCTGGAACTTCGGG | CAGTAGGAAGCGACAAACCC |
| *dgat-2* | TGTGAAGCAAGTGTTCAAAGG | TCATACCGATGCTGAGGAGGA |
| *acl-1* | TGGAATGTGCTTGATTATCG | ATTCGGAAGTGGTTGTTTGG |
| *acl-4* | TGCATCATTCTGGCAACATT | GAGCACCCCACTCGAATATG |
| *lipl-1* | TCCATCAACTGTTGTGCAAAAC | AGGAACCGTTCAAAACATCGT |
| *lipl-3* | CCAGAGTACGACTTTACCGC | GTTGTTCTGCGCAATTATAGCA |
| *lipl-4* | AAAACAAGACCTGGAAGAAACG | ATAAACTTGGCTGGCTGCAT |
| *sbp-1* | GCACGTTCTGACATGTGGAA | CCGCCAAACCCAGATTGTCT |
